# Specialization into Host Sea Anemones Impacted Clownfish Demographic Responses to Pleistocene Sea Level Changes

**DOI:** 10.1101/2024.07.12.603135

**Authors:** Alberto García Jiménez, Théo Gaboriau, Lucy M. Fitzgerald, Sara Heim, Anna Marcionetti, Sarah Schmid, Joris Bertrand, Glenn Litsios, Abigail Shaughnessy, Carl Santiago, Ploypallin Rangseethampanya, Phurinat Ruttanachuchote, Wiphawan Aunkhongthong, Sittiporn Pengsakun, Makamas Sutthacheep, Bruno Frédérich, Fabio Cortesi, Thamasak Yemin, Nicolas Salamin

## Abstract

Fluctuating sea levels during the Pleistocene led to habitat loss and fragmentation, impacting the evolutionary trajectories of reef fishes. Species with specialized ecological requirements or habitat preferences, like clownfishes (Amphiprioninae), may have been particularly vulnerable due to their intricate dependence on sea anemones. The diverse host specializations within this group likely contributed distinct responses to sea-level fluctuations, differentially shaping their recent evolutionary histories. Leveraging a comprehensive genomic dataset, we reveal demographic patterns and connectivity dynamics across multiple populations of ten clownfish species under different host specializations. Host-generalist species demonstrated strong resilience to habitat perturbations, while those specialized on single hosts suffered dramatic bottlenecks linked to sea-level fluctuations. Spatial analyses revealed the significant role of oceanic currents in shaping clownfish genetic diversity landscapes. Dispersal barriers were driven by environmental variables, with the Coral Triangle emerging as a hub of genetic diversity. Our results reveal how clownfish associative behavior influences their population dynamics, holding major implications for their conservation such as the need to consider their mutualism with sea anemones, particularly on host-specialists, to ensure their survival in the face of climate threats. These findings extend broader principles of conservation, improving our understanding of species’ responses to ecological constraints and environmental changes over evolutionary timescales.

## Introduction

Habitat loss and fragmentation pose significant threats to species biodiversity and population resilience across ecosystems, disrupting natural ^1*^habitats and connectivity essential for species persistence [1]. While habitat loss negatively affects species biodiversity and population resilience [2, 3], habitat fragmentation effects are more complex and context-dependent [4, 5]. Habitat fragmentation can promote allopatric speciation and trigger diversification processes [6, 7], while limiting gene flow between previously connected populations, leading to loss of genetic diversity and compromised resilience of populations [8, 9]. The outcome depends on factors such as species’ dispersal abilities, ecological traits, and habitat fragment configuration [10].

Pleistocene sea-level fluctuations caused a global fragmentation of coral reef habitats [11], resulting in habitat loss and increased reproductive isolation for reef species [12]. These changes are hypothesized to have caused significant population bottlenecks and genetic structuring among various reef fish species [13]. However, the impact of limited dispersal and strong ecological adaptations, such as habitat specialization, remains unexplored. This underscores the ongoing need to improve our understanding of how species with different ecological specializations respond to climate change, both historically and in future scenarios, crucial for developing more effective conservation strategies [14].

The interplay between habitat dynamics and species’ ecological adaptations is particularly evident in reef fishes, with clownfishes serving as a compelling example. Clownfishes are known for their unique and tight relationship with sea anemones, which has made them a focal point of marine evolutionary research [15]. This mutualistic relationship is crucial for clownfish survival and reproduction [16], with no observations of clownfishes living independently of sea anemones [17, 18]. Clownfish ecological adaptations significantly impact their population dynamics and vulnerability to habitat changes [19]. Their strong attachment to their host anemones [16], reflects their heavy reliance on the coral reef ecosystem. This dependency is further emphasized by their relatively short pelagic larval dispersal stage of approximately 15 days [20, 21], with larvae rarely dispersing beyond 27 km [22]. Additionally, clownfishes have a unique social structure where only the two largest individuals within a social group form the breeding pair [23]. These factors make clownfish populations particularly sensitive to habitat disturbances, with their mutualistic relationship playing a crucial role in shaping their biogeography and population dynamics [24, 25].

Clownfish species exhibit a generalist-specialist continuum in host usage, reflecting a complex interplay between ecological adaptation and habitat preference. This gradient is primarily influenced by the habitat preferences of their associated sea anemones, indicating an ecological sorting along environmental gradients [26]. Specialized species form tight associations with *Radianthus magnifica, Entacmaea quadricolor*, or members of the *Stichodactyla* genus, while several clownfish species display generalist behavior, capable of inhabiting, socializing, and reproducing across different sea anemone species without clear preference [27]. Variations in sea anemone characteristics create a heterogeneous landscape of resources and protection for clownfish populations [28], likely influencing their ecological and evolutionary trajectories. Despite extensive research on clownfish ecology, the long-term evolutionary impacts of historical habitat changes on their population dynamics remain poorly understood. Quaternary climate fluctuations significantly affected reef communities, but the extent to which clownfishes’ unique host associations may have buffered against these environmental changes is unclear. This knowledge gap highlights the need to investigate how clownfish-anemone mutualism has influenced resilience to environmental shifts over time, providing crucial insights for conservation strategies in the face of ongoing climate change and habitat degradation.

We hypothesize that clownfish populations have undergone distinct evolutionary trajectories shaped by their specialized relationships with sea anemone hosts and Pleistocene sea-level fluctuations. To test this, we analyze 382 geo-referenced genomes from ten clownfish species with varying host specializations. Our multifaceted approach investigates the impact of late Pleistocene sea-level fluctuations on clownfish demographic histories using Sequential Markovian Coalescent methods, aiming to reconstruct historical population size changes and identify potential bottlenecks or expansions. Through spatial genomic analyses, we explore the influence of environmental factors on genetic diversity and dispersal patterns, disentangling the contributions of geographical barriers, habitat preferences, and historical climate changes to current population structures. We also examine differences in demographic patterns and connectivity dynamics across host specializations, assessing the influence of host specificity on long-term population dynamics and evolutionary trajectories.

## Results

We generated whole-genome sequencing data from 382 geo-referenced samples to analyze the population structure and reconstruct the demographic histories of multiple populations across ten clownfish species with different host-specialization regimes (Figure 1). The sequencing reads were mapped onto the *Amphiprion clarkii* reference genome, with mapping rates ranging from 81.05% to 97.74% (Supplementary Figure S1a). Average sequencing depths varied between 5.82X and 42.21X (Supplementary Figure S1b). Generated species-specific SNP datasets contained between 202,007 and 2,192,819 SNPs (detailed information in Supplementary Material).

**Fig 1:**
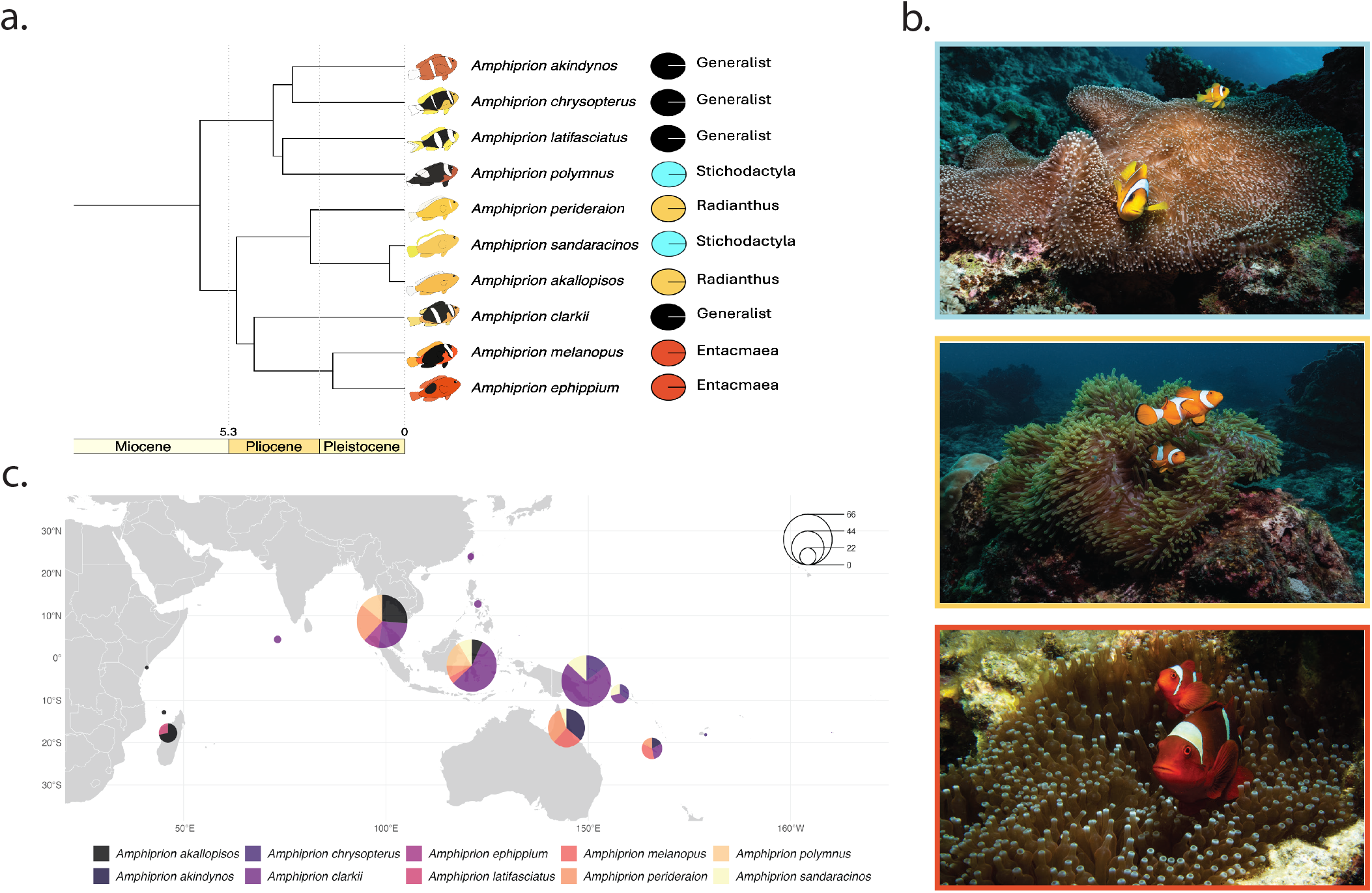
Species in this study. a) Subset of phylogenetic relationships from [27], showing studied species and host specialization types (colored circles). b) Sea anemone exemplars for each host specialization: *Stichodactyla mertensii* (light blue), *Radianthus magnifica* (orange), and *Entacmaea quadricolor* (red). Credit: Lucy Fitzgerald. c) Sample distribution by species. Pie charts show species proportions at each location; size indicates total sample number (see legend). Colors represent species (indicated below).

### Population structure

Phylogenetic relationships within each species were inferred with a GTR + Γ model and 100 bootstraps. These relationships were primarily structured according to geographic locations (Supplementary Figure S3). Some individuals showed greater genetic similarity to those from nearby populations than to con-specifics from their own population, suggesting localized gene flow or potential recent migration events between adjacent populations. Principal Component Analysis supported these findings, clustering individuals by geographic populations. Admixture analyses revealed limited genetic population structure, with the optimal number of genetic clusters (K) consistently lower than the number of geographically defined populations (Supplementary Figure S4). Specialist species showed higher genetic structuring relative to their sampled populations compared to generalists (Supplementary Table S2), highlighting the influence of ecological specialization on the genetic structure of their populations.

### Patterns of genetic diversity

We used the Estimated Effective Migration Surfaces *eems* [29] to analyze spatial patterns of genetic diversity and gene flow across species. The method uses a Bayesian approach to estimate posterior probabilities of genetic diversity and migration rates from pairwise genetic dissimilarities in geo-referenced genomic data, allowing for the identification of areas with significantly higher or lower genetic diversity than expected under an isolationby-distance model.

The spatial distribution of genetic diversity varied among species and regions. Widely distributed generalists,

*A. clarkii* and *A. chrysopterus*, exhibited extensive areas of high genetic differentiation within their populations (Figure 2). These species, along with *A. melanopus* and *A. perideraion* specialists, displayed higher-than-expected genetic diversity within the Coral Triangle, contrasting with predictions based on Isolation-by-distance (*P*_log(*q*)*>*0_ *>* 0.9; Supplementary Figure S5a,c,d and f). In contrast, specialist species, such as *A. akallopisos* in the West Indian Ocean, *A. perideraion* in the Great Barrier Reef, and *Stichodactyla* specialists *A. polymnus* and *A. sandaracinos* in the Coral Triangle, predominantly exhibited lower levels of genetic diversity than expected (*P* (log(*q*) *<* 0) *>* 0.9; Supplementary Figure S5e, f, g and h).

**Fig 2:**
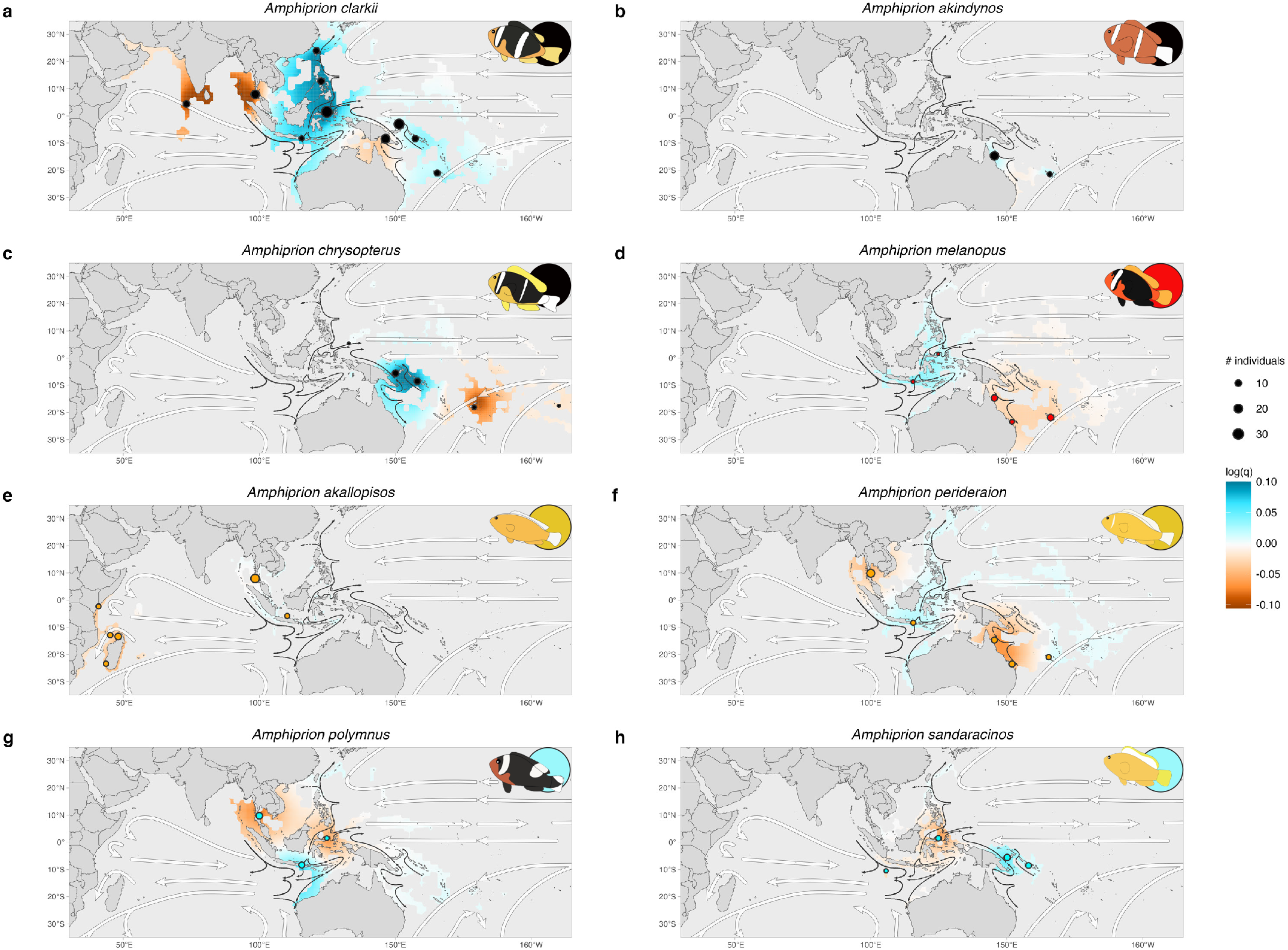
Spatial patterns of genetic diversity. Maps show the clownfish clade distribution range. Sampling locations are indicated by circles, with size representing the number of individuals, and color the host category (black-generalists, orange-*Radianthus*, red-*Entacmaea*, blue-*Stichodactyla*). Effective genetic diversity surfaces are masked by species distributions. White arrows indicate global oceanic currents, while dark arrows indicate Coral Triangle currents. Values reflect log-transformed genetic diversity rates (blue-higher, red-lower than expected under Isolation-by-Distance). Species names are shown at the top, and vector illustrations and host category on the top-right corner.

Indian Ocean populations, including *A. clarkii* in the Maldives and Thailand, and *A. akallopisos* in Madagascar, Mayotte, and Kenya, displayed low genetic differentiation compared to expectations under Isolation-by-distance (*P*_log(*q*)*<*0_ *>* 0.9; Supplementary Figure S5a and e). Similar patterns were observed in the Pacific region, such as *A. chrysopterus* in East Melanesia and Polynesia, and *A. melanopus* and *A. perideraion* in the Great Barrier Reef (*P*_log(*q*)*<*0_ *>* 0.9; Supplementary Figure S5d, and f). The Gulf of Thailand also exhibited lower genetic diversity than expected in *A. perideraion* and *A. polymnus* populations (*P*_log(*q*)*<*0_ *>* 0.9; Supplementary Figure S5f, and g).

This analysis revealed diverse spatial patterns of genetic diversity and gene flow across species, highlighting distinct genetic clusters in specialist and generalist species within the Coral Triangle and Indian Ocean regions. We conducted spatial auto-regressive models to test the explanatory power of oceanographic, temperature, nutrient, and water quality variables, along with clownfish and sea anemone species richness, on genetic diversity patterns. Combined explanatory variables better explained genetic diversity distribution across species than single environmental variables, underscoring the complex interplay shaping population genetic structures in clownfish. Best models included temperature, salinity, and current velocity, while ecological variables, such as species diversity of clownfish and sea anemone, had limited influence (Supplementary Table S4).

### Population connectivity and dispersal barriers

*eems* analyses inferred higher migration rates among demes within the Coral Triangle than expected under a stepping-stone model for multiple species (Figure 3), with notable rates observed in *A. clarkii, A. perideraion*, and *A. polymnus* (*P*_log(*m*)*>*0_ *>* 0.9; Supplementary Figure S6a, f and g). Lower migration rates were identified in regions like Melanesia, the Gulf of Thailand, the Sunda Shelf, the Sahul Shelf, and the area connecting Thailand and East African populations of *A. akallopisos* (*P*_log(*m*)*<*0_ *>* 0.9; Supplementary Figure S6).

**Fig 3:**
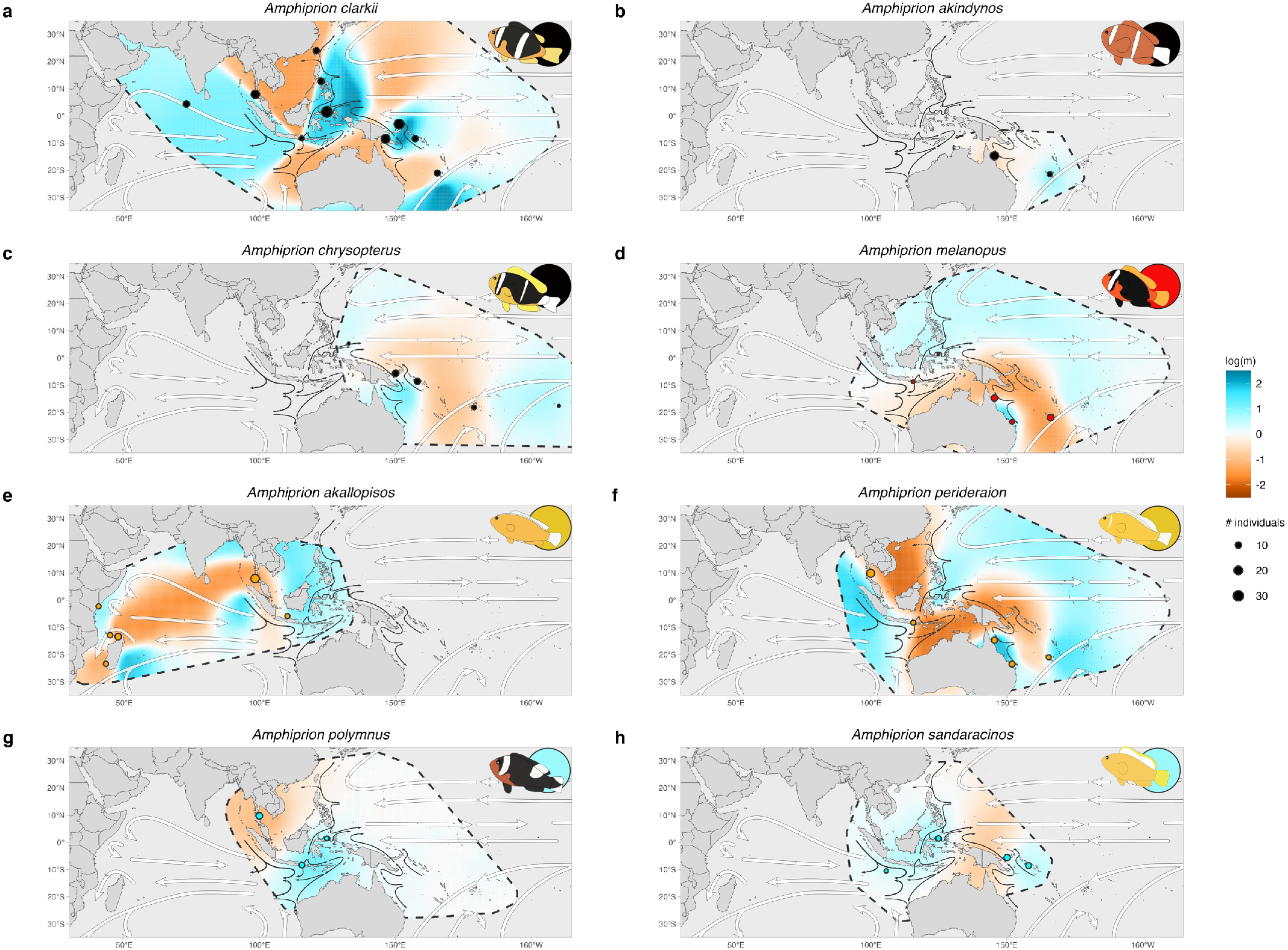
Spatial patterns of population connectivity. Maps show the clownfish clade distribution range. Sampling locations are indicated by circles, with size representing the number of individuals, and color the host category (black-generalists, orange-*Radianthus*, red-*Entacmaea*, blue-*Stichodactyla*). White arrows indicate global oceanic currents, while dark arrows indicate Coral Triangle currents. Values reflect log-transformed effective migration rates (blue-higher, red-lower than expected under a stepping-stone process). Species names are shown at the top, and vector illustrations and host category on the top-right corner.

Migration patterns differed between generalist and specialist species. The most widespread and generalist species, *A. clarkii*, displayed the highest migration rates in the Coral Triangle, with values two orders of magnitude higher than those expected under a stepping-stone model (Supplementary Figure 3a). In contrast, specialists exhibited reduced migration rates, with *A. melanopus, A. akallopisos* and *A. perideraion* displaying migration rates from 30 to 80 times lower than expected under a steppingstone process (Figure 3d, e, and f).

Migration rates among clownfish species in the Coral Triangle exceeded expectations based on Isolation-by-distance, while shallow continental shelves and vast bodies of water posed significant geographical barriers to gene flow. The inferred migration patterns are closely linked to the Indonesian Through-Flow and prevailing oceanic currents. Analogous to spatial analysis of genetic diversity patterns, spatial auto-regressive models revealed ocean currents, temperature and water quality as the main variables explaining effective migration rates across most clownfish species. Nevertheless, the most comprehensive model, integrating physical, chemical, and ecological variables, frequently outperformed others, as indicated by the lowest corrected Akaike Information Criterion (Supplementary Table S5).

### Demographic history reconstruction

We employed *msmc2* (Multiple Sequentially Markovian Coalescent; [30] to infer historical changes in effective population size (*N*_*e*_) from whole-genome sequences. To ensure the robustness of our reconstructions, we generated ten independent *msmc2* runs per species per population, each using three randomly selected individuals.

*N*_*e*_ reconstruction through time revealed distinct demographic patterns between specialist and generalist clownfish species during the Quaternary period. Specialists species exhibited a decrease in *N*_*e*_, regardless of their host association (Figure 4b). This declines coincided with the trend of decreasing sea level during the late Pleistocene, reaching a minimum approximately 20,000 years ago (Figure 4a). Notably, specialist species (*Entacmaea, Radianthus*, and *Stichodactyla* specialists) initially exhibited higher historical *N*_*e*_ than generalists (Figure 4c), with peak values observed between 1 million and 200,000 years ago.

**Fig 4:**
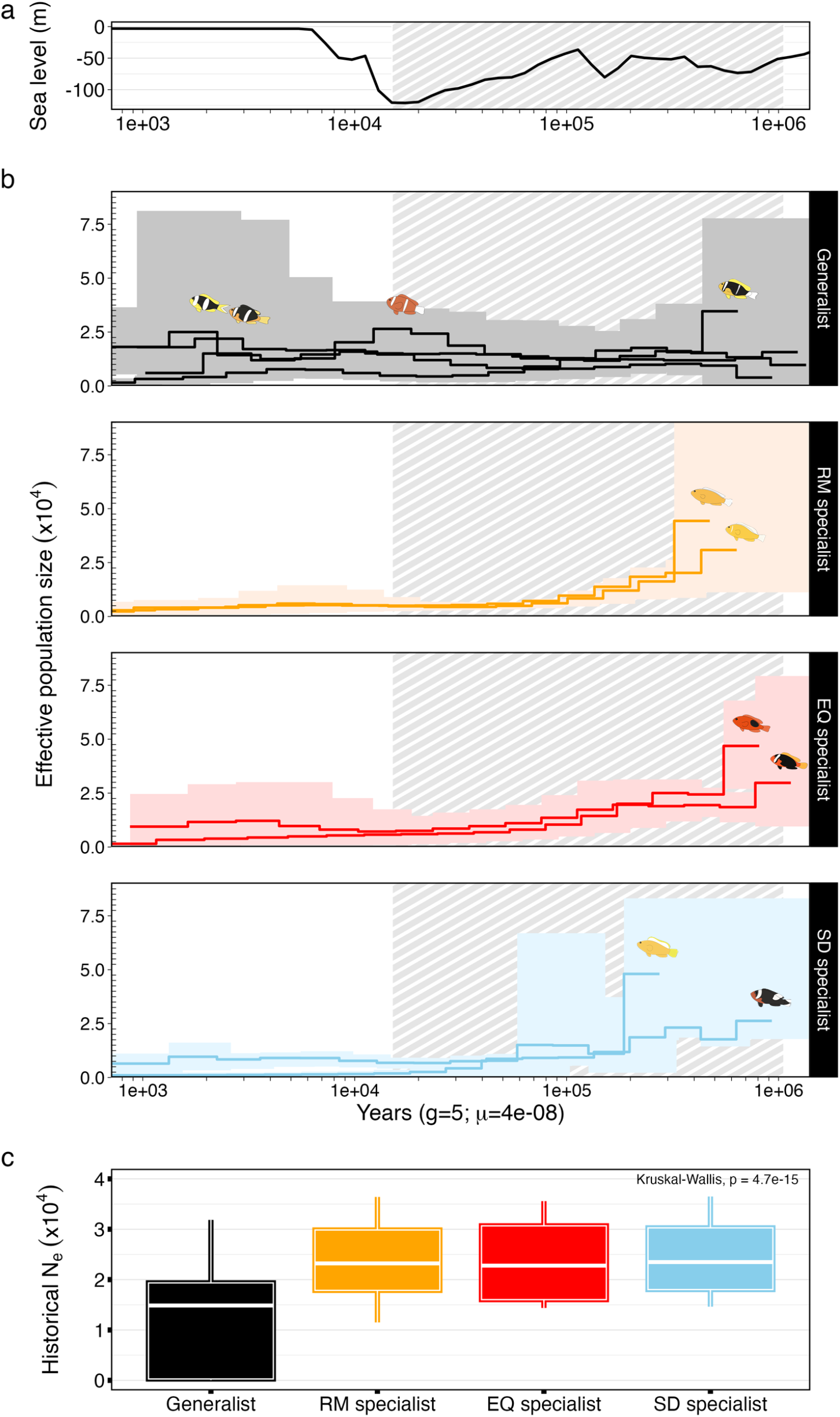
Comparison of demographic reconstructions among host specialization groups in the context of late Pleistocene sea level fluctuations. (a) Sea level dynamics from 1 million years ago to present, with declines indicated by striped grey lines. (b) Demographic history reconstruction using *msmc2* [30] for ten species. Panels show different host associations: generalists (black), *Radianthus* (orange), *Entacmaea* (red), and *Stichodactyla* (blue). Step-lines represent average demographic reconstructions per species, with species illustrations near the highest *N*_*e*_ values. Light-colored areas show the range of *N*_*e*_ values across all populations. Period of sea-level declines is highlighted by striped grey lines. (c) Historical *N*_*e*_ for each group, calculated as the weighted harmonic mean of *N*_*e*_ estimates. The white line is the median, colored boxes show the IQR, and whiskers denote bounds. Kruskal-Wallis significance is indicated at the top right.

While specialist species showed significant declines in *N*_*e*_ estimates over time (*Radianthus* specialists: W = 2278; *p-value* < 0.001; *Entacmaea* specialists: W = 55; *p-value* < 0.001; and *Stichodactyla* specialists: W = 496; *p-value* < 0.001), generalists maintained relatively stable *N*_*e*_ with no significant variations across the periods of sea level change (W = 3665; *p-value* = 0.679; Figure 5a and Supplementary Figure S8a). Generalist species showed an upward trend of *N*_*e*_ starting around 100,000 years ago, with the exception of *A. chrysopterus*. Moreover, *N*_*e*_ dynamics in generalist species appeared more variable, displaying species-specific fluctuations over time compared to the consistent declining trends observed in all specialist species.

**Fig 5:**
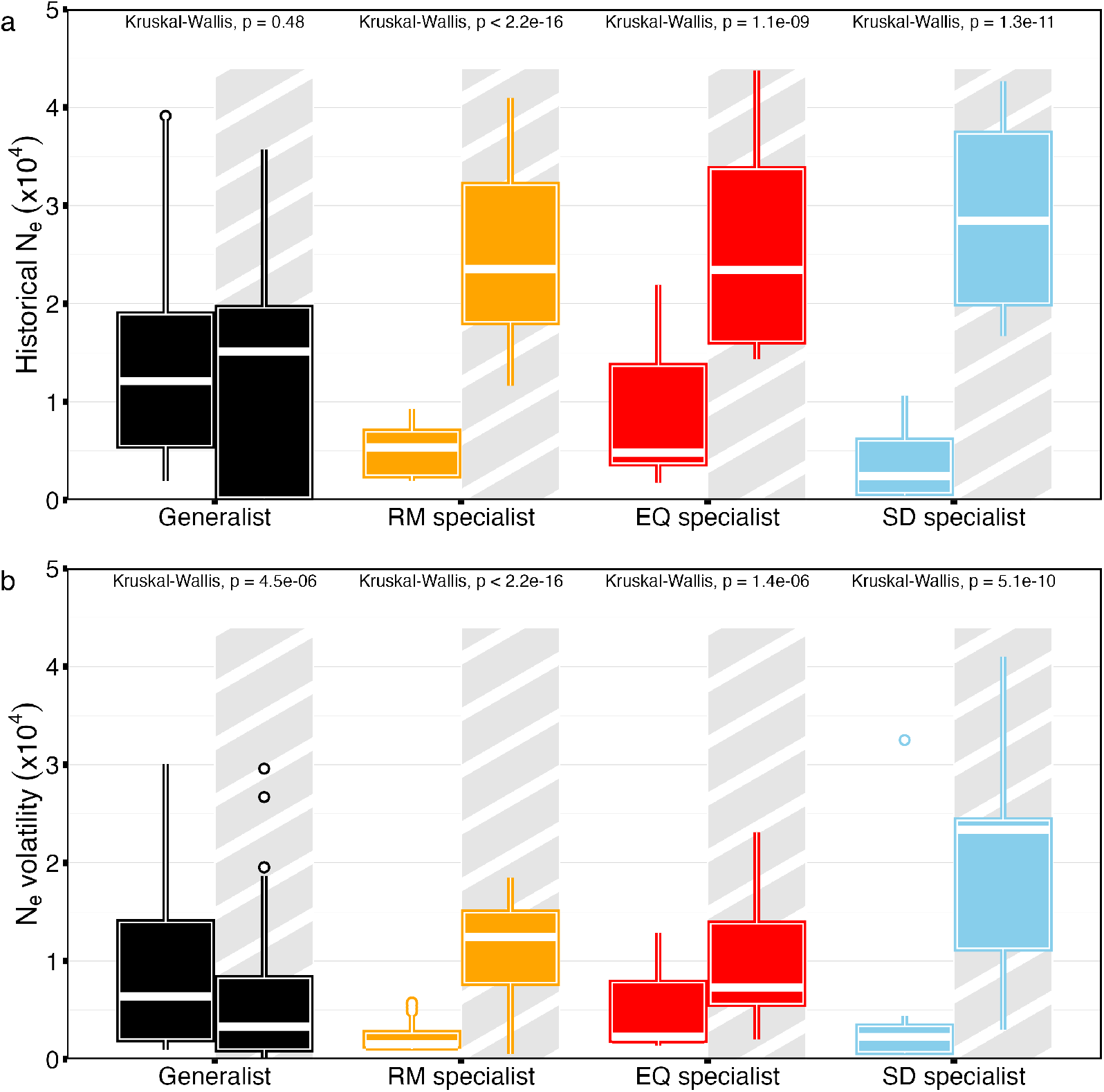
Comparisons of *N*_*e*_ between the period of sea level declines and subsequent rise across host specialization groups. Values relative to the period of sea-level declines are differentiated by a striped grey-shadowed background. White line represents the median while colored boxes indicate the IQR. Whiskers denote the upper and lower bounds. KruskalWallis statistical significance for each comparison is displayed at the top of each box plot. a) Historical *N*_*e*_, calculated as the weighted harmonic mean of *N*_*e*_ estimates over time for each population’s *msmc2* reconstruction. b) *N*_*e*_ volatility, calculated as the average true range (ATR) over 10,000-year intervals.

The most recent *N*_*e*_ estimates, within the last 1,000 years, revealed critically low values for all specialist species. The most extreme case was observed in the Papua New Guinea population of *A. sandaracinos*, where as few as 177 *±* 146 individuals were estimated to contribute to the next generation approximately 180 years ago. An exception is the Indonesian population of *A. melanopus*, with relatively high *N*_*e*_ of 29, 182*±* 2.49 individuals around 3,000 years ago, comparable to the *N*_*e*_ values of the generalist *A. clarkii* in Indonesia during the same period (Supplementary Figure S11).

Average *N*_*e*_ volatility across 10,000-year periods, a measure of demographic fluctuation intensity, was initially similar across all groups (Figure S9), ranging between 1,250 and 2,500 individuals contributing to the genetic diversity pool. However, following the period of sea level declines, all specialist species exhibited a significant decrease in *N*_*e*_ volatility (*Radianthus* specialists: W = 7, *p-value* < 0.001; *Entacmaea* specialists: W = 158, *p-value* < 0.001; and *Stichodactyla* specialists: W = 10, *p-value* < 0.001). In contrast, populations of generalist species showed significant increases in *N*_*e*_ over time after this period (W = 4678; *p-value* < 0.01; Figure 5b).

## Discussion

Our demographic reconstructions and spatial analyses of clownfish populations across the Indo-Pacific reveal a complex interplay of historical, ecological, and oceanographic factors shaping their current distribution and genetic structure. The findings underscore the significant impact of Pleistocene sea-level fluctuations, host specialization, and oceanic currents on clownfish populations.

### Historical Influences on Population Demography

Previous studies have hypothesized that sea-level fluctuations during the Pleistocene drove global population bottlenecks and structured populations across various marine taxa [12]. During lower sea levels, marine habitats fragmented, leading to population isolation and promoting genetic differentiation [31, 32]. As sea levels rose, previously isolated populations reconnected, facilitating gene flow and population mixing. These dynamic processes have contributed to genetic bottlenecks [13], where populations undergo drastic reductions in size, followed by expansion as environmental conditions improve.

Our estimated historical effective population sizes provide evidence of the impact of sea-level fluctuations on clownfish demographic dynamics. Specialist species— *Radianthus, Entacamea*, and *Stichodactyla* specialists— experienced significant declines in *N*_*e*_, leading to population bottlenecks following the sea-level drop between 500,000 and 20,000 years ago, without subsequent recovery during rising sea levels. This highlights the long-lasting consequences of these bottlenecks and underscores the critical interplay between host specializations and environmental changes. In contrast, generalist species exhibited greater population stability, likely due to their ability to exploit diverse hosts, mitigating the severe consequences of habitat loss and fragmentation caused by sea-level fluctuations during this period.

### Ecological and Oceanographic Factors

While sea-level declines during the Pleistocene likely affected host sea anemones similarly [among other marine organisms 12], clownfish population dynamics might not have entirely overlapped with their hosts. Specialists may have followed the fate of their singular host sea anemones, experiencing similar declines due to habitat loss and fragmentation [33]. Conversely, generalists, with their capacity to exploit diverse hosts, likely had higher survival chances during sea-level declines [34, 35]. An upward trend in effective population size of generalist populations around 100,000 years ago could be attributed to their opportunistic behavior, which allowed them to exploit habitats vacated by specialists undergoing bottlenecks. This phenomenon, observed in other taxa in response to climate change [36], is supported by fossil records indicating that past ecosystems shifted towards dominance by generalist species during such periods [37].

Clownfish exhibit a strong attachment to their host anemones [16], coupled with short pelagic larval dispersal and robust social and reproductive structures, rendering them highly vulnerable to habitat disturbances [19]. During periods of sea level rise, oceanographic conditions would have played a pivotal role in shaping the regional recolonization dynamics of clownfish, akin to their influence on other marine species [38, 39]. These oceanographic factors likely underlie the distinctive genetic diversity and population structure observed across clownfish species. Our findings strongly indicate that Pleistocene sea-level fluctuations and associated oceanographic dynamics have profoundly influenced the present biogeography of clownfish.

These dynamics likely contributed to the allopatric speciation events within the clownfish clade during this epoch [40, 41]. The synchronous occurrence of diversification bursts and significant bottleneck events suggests that habitat fragmentation could have facilitated these speciation processes. In many other taxa, habitat fragmentation due to climatic changes or natural barriers has similarly driven allopatric speciation and diversification [e.g., 6, 7]. In the case of clownfish, Pleistocene sea-level fluctuations likely created isolated populations, potentially accelerating speciation through allopatric mechanisms. However, we cannot disregard the importance of their unique social structure and short larval dispersal in driving speciation [42], particularly after geographical expansions of the late Pliocene and early Pleistocene [27].

### Dispersal Barriers and Connectivity

Genetic connectivity analyses identified key geographical barriers to clownfish dispersal, including the Gulf of Thailand, Melanesia, and the Indian Ocean Gyre. These barriers, shaped by tidal and oceanic currents, contribute to genetic isolation and differentiation among populations. Our results highlight the profound impact of oceanic currents on clownfish dispersal routes, consistent with findings in other marine taxa [43–45]. The Indonesian Through-Flow plays a vital role in facilitating genetic exchange and increasing genetic diversity among clownfish populations by linking the Indian and Pacific Oceans. Conversely, areas at the margins of clownfish distributions, such as the Western Indian Ocean and Polynesia, exhibit the lowest levels of genetic differentiation due to dispersal barriers like the Indian Ocean Gyre and the Sunda and Sahul shelves.

Integrating these findings with demographic reconstructions provides insight into how habitat fragmentation during Pleistocene sea-level declines induced reproductive isolation. This process likely restricted gene flow between previously connected populations, resulting in genetic diversity loss and compromised population resilience [8, 9, 13]. Despite reconnecting during sealevel rises, tidal and current dynamics in certain areas have perpetuated isolation among populations. This underscores the crucial role of ocean currents in mediating the dispersal of fish larvae between fragmented habitats [a notion suggested by 46–48] and highlights the importance of incorporating dispersal paths in marine conservation efforts.

### The Coral Triangle as a Biodiversity Hotspot

The Coral Triangle, known for its unparalleled species richness and endemic diversity [49], boasts an abundance and complexity of reef habitats. Combined with the Indonesian Through-flow that links the Indian and Pacific Oceans, these factors have been crucial in shaping the current marine biodiversity patterns in the Indo-Pacific. This region is particularly pivotal for clownfish, representing the center of origin for the clade [40] and currently hosting the highest diversity of clownfish species [50, 51].

Beyond generating biodiversity, our findings underscore the crucial role of the Coral Triangle in maintaining genetic diversity within species. Populations in this region exhibit higher historical *N*_*e*_ values than those in other regions, aligning with its historical role as a critical refuge during Pleistocene sea-level fluctuations [11]. The ecological complexity and environmental heterogeneity of the reef ecosystems in the Coral Triangle have likely contributed to maintaining these higher levels of genetic diversity [52–54].

## Conclusion

Our study provides crucial insights into the dynamics of clownfish populations through an integrated analysis of Pleistocene sea-level fluctuations, population dynamics, and genetic diversity across their range. We highlight how ecological specializations influence species’ responses and adaptability to climate change and environmental fluctuations. Our comprehensive approach identifies regions prone to increased reproductive isolation and inbreeding, alongside areas that serve as genetic diversity reservoirs. By examining how historical climate events have shaped the current genetic landscape of clownfish species, we can better predict the impacts of ongoing and future climate change on different populations, considering their host specialization and geographic distribution.

These findings are pivotal for guiding conservation efforts, emphasizing the need to prioritize strategies that uphold genetic diversity and ecosystem resilience throughout the clownfish’s habitat. Incorporating historical demographic analysis into conservation planning enhances our understanding of long-term processes affecting clownfish populations, enabling the development of effective strategies to preserve their genetic diversity and adaptive capacity in the face of ongoing environmental changes. Specifically, conservation initiatives should prioritize maintaining connectivity in historical corridors for gene flow, such as the Indonesian Through-Flow, while safeguarding isolated populations that may harbor unique genetic adaptations. This holistic approach will sustain the adaptive potential of clownfish populations amidst habitat degradation and climate change, crucial for the enduring survival of these iconic coral reef inhabitants.

Future research should integrate these findings with contemporary ecological data and direct observations to offer a comprehensive understanding of clownfish population dynamics. Additionally, investigating the current and potential future impacts of climate change on clownfish and their host anemones is imperative for predicting and mitigating challenges facing these emblematic reef fishes.

## Methods

### Sampling and Data Collection

A total of 382 fin clips were collected from 10 clownfish species, representing distinct host categories within the clade as classified by [27]. The samples were systematically collected by capturing fish within their host anemones during SCUBA diving expeditions (see Table S1). Samples were preserved either in 70% ethanol (ETOH) or DESS buffer [55] and stored at 6*°*C immediately after retrieval. Collection efforts spanned multiple locations over an 11-year period starting in 2012 (see Supplementary Material).

### Host Association

Species were categorized based on their host preferences following [27]. Two species, *Amphiprion akallopisos* and *A. perideraion*, classified as RM specialists, have adapted to inhabit and reproduce within *Radianthus magnifica*, a solitary sea anemone species with moderate-length tentacles found in coral reef habitats with moderate water flow. *A. melanopus* and *A. ephippium*, classified as EQ specialists, associate with *Entacmaea quadricolor*, a colonial sea anemone with long tentacles inhabiting crevices and rocky substrates in shallow coral reefs. Finally, *A. sandaracinos* specialized in *Stichodactyla mertensii* and *A. polymnus* specialized in *Stichodactyla haddoni* are classified as SD specialists, as both species belong to the *Stichodactyla* genus, characterized by short-tentacle solitary anemones forming a carpet-like extension over sandy areas of coral reefs. The remaining four species examined (*A. latifasciatus, A. akindynos, A. chrysopterus*, and *A. clarkii*) were categorized as Generalists, showing no clear preferences for any specific host sea anemone species.

### Environmental and Geographical Data Collection

Sea level dynamics over the last million years were obtained from [56], and current climate data were sourced from GMED [57]. Environmental variables included current velocity, surface current, tide average, temperature, bottom temperature, sea surface temperature range, chlorophyll concentration, phytoplankton concentration, nitrate concentration, salinity, dissolved oxygen range, and bottom dissolved oxygen. Environmental data were standardized to a resolution of 0.083*°* x 0.083*°* equivalent to a grid size of 100 km^2^ near the equator, and adjusted to the Indo-Pacific study area. Geographically defined marine regions were obtained from Marine Ecoregions Of the World [MEOW; 58] and shallow reef habitats locations in the Indo-Pacific Ocean from UNEP-WCMC [59] to implement geographical constrains into species distribution models (SDMs). Oceanic current data and Indonesian Through-Flow (ITF) polygons were derived from [60] and [61], respectively, for visualization and interpretation alongside genomic results.

### Species Distributions

We estimated distributions of clownfish and their host sea anemones using 3,340 occurrences of 28 clownfish species (average 115 *±* 150 per species) and 1,211 occurrences of the 10 sea anemones species hosting clownfishes (average 121 *±* 58 per species) sourced from GBIF [62]. Species distribution models were conducted using the *ENMTML* R package [63] to delineate the geographic ranges and distributions of each species. See Supplementary Material for details on data and modeling parameters. Additionally, we inferred clownfish and sea anemone species richness distributions by aggregating presence/absence distribution maps across all clownfishes and all sea anemones, respectively.

### Molecular Data Collection

#### DNA extraction, library preparation and sequencing

DNA was extracted using DNeasy blood and Tissue Kit (Qiagen GmbH, Hilden, Germany) and quantified using Qubit^®^ 2.0 Fluorometer (Thermo Fisher Scientific, Waltham, USA). The integrity of the samples was assessed by electrophoresis. Libraries for samples collected before 2021 were prepared from 100 ng of DNA using the TruSeq Nano DNA library prep standard protocol, while libraries from samples collected after 2021 were prepared using the Nextera DNA Flex Library Preparation Kit following manufacturer’s instructions. Fragment length distribution of the libraries was validated using a Bioanalyzer (Agilent Technologies, Santa Clara, USA). Libraries generated in 2017 and 2018 were sequenced on Illumina HiSeq 2500 platform with 100 paired-end lanes, while libraries prepared in 2019 were sequenced on the Illumina 4000 HiSeq platform with 150 paired-end lanes. Libraries prepared in 2022 and 2023 were sequenced on the NovaSeq 6000 Sequencing System (Illumina, San Diego, California) with 150 paired-end lanes. Sequencing was performed at the Genomic Technologies Facility (GTF) of the University of Lausanne, Switzerland.

#### Reads processing, mapping and SNP calling

Low-quality raw reads were trimmed, and potential adapter contamination was removed using *Trimmomatic* (parameters:–qual-threshold 20 –length-threshold 50, v.0.39; [64]). The quality of the processed reads was assessed with *FastQC* v.0.11.9 [65] and *MultiQC* [66]. Reads were mapped against *Amphiprion clarkii* reference genome (GenBank; ID: JALBFV000000000; [67]) using *BWA* v0.7.17 [68] and *Samtools* v1.15.1 [69] was used to process and sort the mapped reads, generating mapping statistics with *Bamtools* v2.5.2 [70]. The mapping results were processed to reduce potential SNPs calling errors [71]. Mapped reads were filtered with *Samtools* v1.15.1 [69] to retain only primary alignments and proper pairs (i.e., at the expected insert size and orientation) with a mapping quality higher than 30. Redundant sequencing data originated from the overlap of paired reads was removed using the `mergeReads` task from *ATLAS* v0.9.9 [72]. We used *ATLAS* pipeline for the SNP calling as it was shown to be more reliable than *GATK* in non-model species, maintaining high accuracy in variant calling for moderately divergent species [73]. Genotype likelihoods (GL) were calculated at each position using the `GLF` task from *ATLAS* for each sample and the reference genome. Then, genotype likelihoods were used to infer the major and the minor alleles for each position running the majorMinor task, obtaining the individuals genotype. SNP calling pipeline was carried out with all individuals from all species together generating an all-species dataset, and for each species independently, using all individuals from multiple populations, generating species-specific datasets.

#### Variant filtering and data processing

Reliability of the SNP dataset was improved by filtering invariant or poor confidence positions with *vcftools* v1.15.1 [69]. We filtered out lowconfidence positions (variant quality score lower than 30), positions with coverage higher than 50, and with more than 10% of missing data (parameters: –minQ 30 –max-missing 0.9 –minDP 5 –max-meanDP 30 –maxDP 50). The VCF files were split by species populations to estimate individuals heterozygosities (–het) and minor allele frequencies (–maf) using *vcftools* v1.15.1 [69]. With this information, we removed rare variants suspected to be private alleles or sequencing errors using a MAF threshold of 0.02. Further, we merged species population VCF files for each species, keeping only common variants using merge task from *bcftools* v1.15.1 [69]. Finally, VCF files were LD pruned and converted to BED files using *Plink* [74].

#### Population Structure Analysis

We assessed species population structure using Admixture [75] and Principal Component Analysis (PCA) with *Plink* v1.9 [74], aiming to identify genetic clustering patterns between and within species. We carried out Admixture and PCA on each species BED file to determine population identity and assess population structure, crucial for Sequential Markov Coalescence model interpretations [76]. Admixture was ran with K from 1 to the number of assumed populations of each species plus three, with 10-fold cross-validation and 2,000 bootstraps. Following [77], the best K value was selected upon the lowest cross-validation error. We also carried out PCA on all-species dataset to determine species identities. Three *A. polymnus* samples were clustering with *A. clarkii* showing to be potentially misidentified or mislabelled. Consequently, these three samples were removed for demographic reconstruction and *eems* analysis. Additionally, we reconstructed phylogenetic relationships of all samples using *PhyML* [78] with a GTR + Γ and 100 bootstraps to explore the evolutionary relationships among species, complementing population structure analyses.

#### Spatial Connectivity and Effective Migration Rates

We first evaluated Isolation-by-distance patterns by fitting a linear regression between geographical distances and genetic distances. Geographical distances were calculated as the geodesic distance between population centroids using the geoDist function from the *oce* R package [79]. We estimated population pairwise Fst for each species using vcftools v1.15.1 [69] to quantify the genetic distances between populations. The positive association between geographical and genetic distances indicated a clear Isolation-by-distance signal, thereby validating the use of the *eems* method [29], which is based on Isolation-by-distance expectations. We used the Estimated Effective Migration Surfaces *eems* [29] to assess genetic connectivity patterns within and between populations. *eems* estimates effective migration rates and genetic diversity using genetic data, modeling spatial gene flow patterns. Employing Markov Chain Monte Carlo (MCMC) simulations, *eems* infers these rates between demes, providing insights into genetic structure, barriers to gene flow, and dispersal dynamics. It also calculates a posterior probability for each estimate, offering a probabilistic assessment of whether rates are higher or lower than expected under the Isolation-by-Distance assumption. This Bayesian approach facilitates a nuanced understanding of genetic connectivity and the impact of spatial factors on population dynamics and evolutionary processes.

Genetic dissimilarities between pairs were computed using the bed2diffs program, available within the *eems* toolkit. All analyses were executed with 2,000,000 iterations following a burn-in phase of 1,000,000 iterations for the Markov Chain Monte Carlo. For each species, we conducted analyses with 50, 200, and 500 demes, employing three chains for each configuration. This extensive exploration allowed us to compare results and fine-tune parameters, following the developers recommendations. The outcomes of these three runs were visualized using the *rEEMSplots* R package [29], demonstrating consistent results across runs. Species *A. latifasciatus* and *A. ephippium* were excluded from these analyses due to having only one population of each.

#### Spatial Statistical Analyses

We examined the relationship between effective migration and genetic diversity rates derived from *eems* analysis and environmental variables. Initially, we gauge the significance of environmental variables in explaining effective migration surfaces within generalized linear models (GLMs). Then, we assessed spatial auto-correlation using Moran’s on each GLM. Upon confirming spatial auto-correlation’s significance, we applied Rao’s score tests using lm.RStests from the *spdep* R package (Bivand et al., 2011) to determine suitable spatially dependent linear models. Further analysis involved fitting Lagrange Multiplier tests for Spatial Autoregressive (SAR) models using lagsarlm from the *spatialreg* R package [80], independently testing each environmental variable alongside clownfish and sea anemone richness. Additionally, we explored various combinations of variables based on biological expectations (see Supplementary Material for detailed model descriptions).

#### Demographic history reconstruction

We employed *msmc2* [30] to reconstruct the historical dynamics of effective population size (*N*_*e*_) across all populations of the ten clownfish species, following guidelines from *msmc-tools* (https://github.com/stschiff/msmc-tools). From individual BAM files, we used `mpileup` and call tasks in *bcftools* v1.15.1 [69] followed by `bamCaller.py` from *msmc-tools* to generate chromosome-level individual VCF files. Average coverage by chromosome was estimated using depth task in *Samtools* v1.15.1 [69]. VCF files were phased with *Whatshap* [81], and consensus and individual masks were generated using `makeMappabilitymask.py`. Then, *msmc2* input files were generated with `generate_multihetsep.py`.

For each species, we conducted ten population reconstructions, each time selecting three random individuals per population. The time-segmented pattern used was 1 *∗*2 + 16 *∗* 1 + 1*∗* 2, recommended for shorter genomes [30], with a maximum of 100 iterations. Additionally, we performed 50 bootstraps using multihetsep_bootstrap.py to assess uncertainty levels at each time step. Historical time periods and *N*_*e*_ were inferred using a fixed per-generation mutation rate of 4*×* 10^−8^ [82, 83] and generation time of 5 years [84], consistent with previous clownfish studies [85, 86].

Population *N*_*e*_ reconstructions for each species were averaged to derive overall species trajectories. Historical *N*_*e*_ values were computed using the weighted harmonic mean, which adjusts for the relative time intervals to account for dataset unevenness. Additionally, we assessed the volatility of species *N*_*e*_ using the Average True Range (ATR), a statistical measure commonly used in financial markets to gauge volatility. *N*_*e*_ volatility was calculated as the average of the true ranges over successive 10,000-year intervals. This metric provides insights into the fluctuation of *N*_*e*_ over short evolutionary timescales, helping to assess the stability or volatility of population sizes across evolutionary periods and enhancing our understanding of population dynamics and their responses to environmental changes.

## Supplementary information

## Acknowledgements

We express our gratitude Rosanna Pescini for the DNA extractions, to the Lausanne Genomic Technologies Facility for their assistance in sequencing the samples and to the DCSR infrastructure of the University of Lausanne for providing computational resources essential for data analysis. Special thanks are extended to the staff at the Lizard Island Research Station for their invaluable support during fieldwork and acknowledge the Dingaal, Ngurrumungu and Thanhil peoples as traditional owners of the lands and waters of the Lizard Island region. We are deeply appreciative of the local authorities of Australia, Indonesia, Thailand, Kenya, Mayotte, Madagascar, Maldives and New Caledonia for granting permits to collect samples and their assistance with field logistics. In particular, we extend our gratitude to the Quesnel family and Victoria from COREsea for their invaluable help and logistical support during fieldwork in New Caledonia and Thailand, respectively. Finally, we thank Daniele Silvestro, Diego Hartasanchez and Thibault Latrille for their useful comments on the manuscript.

## Declarations

### Funding

Financial support for this research was provided by the University of Lausanne funds and the Swiss National Science Foundation (Grant Number: 310030_185223). We also acknowledge support from the Australian Research Council (ARC) with the Discovery Early Career Research Award DE200100620 and Discovery Project DP180102363 to FC, contributing to this study.

### Conflict of interest statement

The authors declare no competing interests in the publication of this work.

### Biological Sampling Permits

The research permits obtained for this study are the following:

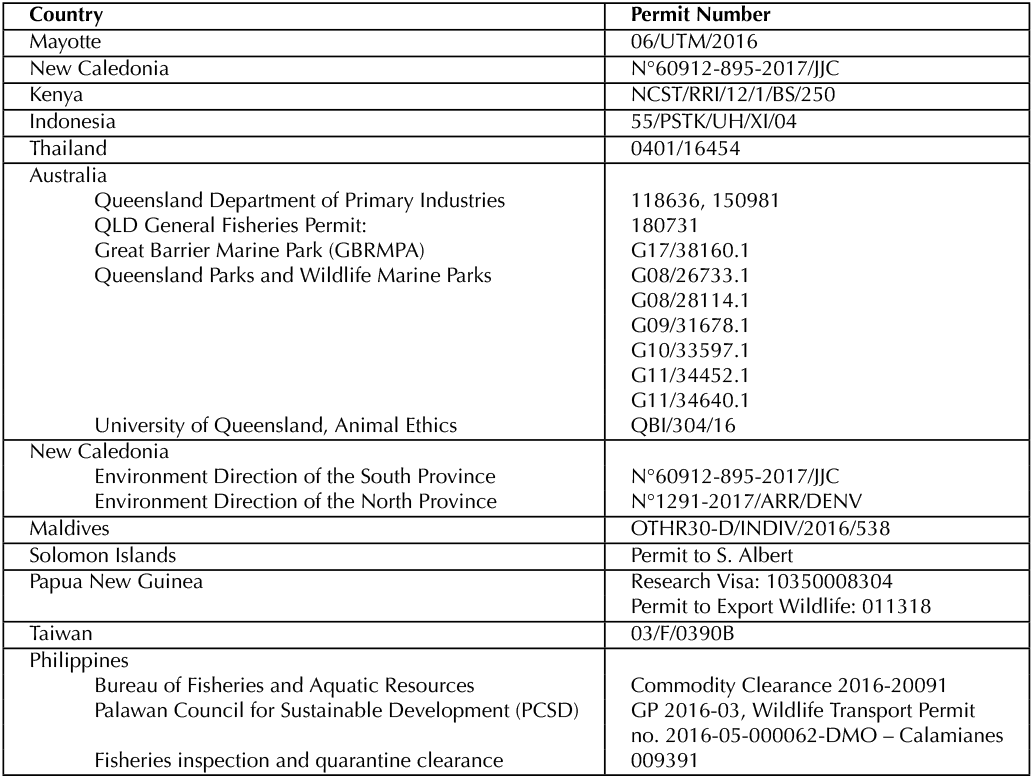

### Benefits Generated

The sharing of our data and results on public databases ensures transparency and facilitates further research in the field. All fieldwork conducted during this study adhered to local regulations and was carried out in collaboration with local entities, including the University of Mayotte (Mayotte), University of New Caledonia (New Caledonia), Kenya Marine and Fisheries Research Institute (Kenya), National Research Institute (Papua New Guinea), Universitas Hasanuddin (Indonesia), University of Ramkhamhaeng (Thailand), University of Toliara and the Ministry of Marine Resources and Fisheries (Madagascar), Lizard Island Research Station and University of Queensland (Australia). Our research activities were conducted in accordance with the “Access and Benefit Sharing” (ABS) principles of the Nagoya Protocol established by the Convention on Biological Diversity (CBD).

Samples from Indonesia (2013), Maldives (2017), New Caledonia (2017), Thailand (2022), Madagascar (2022) and Australia (2023) were obtained by Nicolas Salamin’s research group. Samples from Solomon Islands, Papua New Guinea and Lizard Island (2018) were obtained by Cynthia Riginos’ research group. Sampling in the Solomon Islands was via the Australian Government’s Pacific Strategy Assistance Program and with the assistance of the Roviana Conservation Foundation. Samples from Taiwan were obtained by Filip Huyghe. Samples from Philippines were obtained by Jimmy O’Donnell.

## Data availability

Metadata associated with our study will be uploaded to GEOME, and genomic data deposited in the National Center for Biotechnology Information (NCBI) under a BioProject ID. Access to these data will also be provided upon acceptance of the manuscript.

## Code availability

The scripts used for the analyses will be deposited on Dryad upon acceptance of the manuscript, ensuring the reproducibility and accessibility of our research findings.

## Author contribution

AGJ and NS designed the study. AGJ, TG, LF, SH, AM, SS, JB, GL, LM, BF, FC, AS, CS, PR, PR, TW, SP and TY participated in the fieldwork expeditions and sample collection. AGJ conducted the research and analyses and wrote the manuscript. TG and NS contributed to the interpretation of the analyses, and the writing of the manuscript. All authors participated in reviewing, correcting, and approving the final version of the manuscript.

## Notes

### Competing Interest Statement

The authors have declared no competing interest.

