## Supplementary Material for "Specialization into Host Sea Anemones Impacted Clownfish Demographic Responses to Pleistocene Sea Level Changes"

#### Supplementary Results

##### All-species PCA

All-species data set was composed of 2,612,217 SNPs after filtering and pruning.

The first two principal components of the PCA on the all-species dataset explained a total of 42% (PC1 28.6% and PC2 13.3%) of genetic variability and differentiated three major genetic clusters: i) all individuals of the most generalist species *A. clarkii*, as well as three *A. polymnus* individuals from Indonesia; ii) individuals from *Entacmaea* (EQ) specialist species *A. ephippium* and *A. melanopus*; iii) the remaining individuals from the species *A. latifasciatus*, *A. akindynos*, *A. chrysopterus*, *A. akallopisos*, *A. perideraion*, *A. sandaracinos*, and *A. polymnus*. Individuals of the last cluster appeared grouped by species, with the exception of *A. akindynos* and *A. chrysopterus* that showed some overlap. One *A. akallopisos* individual also appeared clustering closer to *A. sandaracinos* group (Figure S2).

#### Supplementary Figures

**a.**

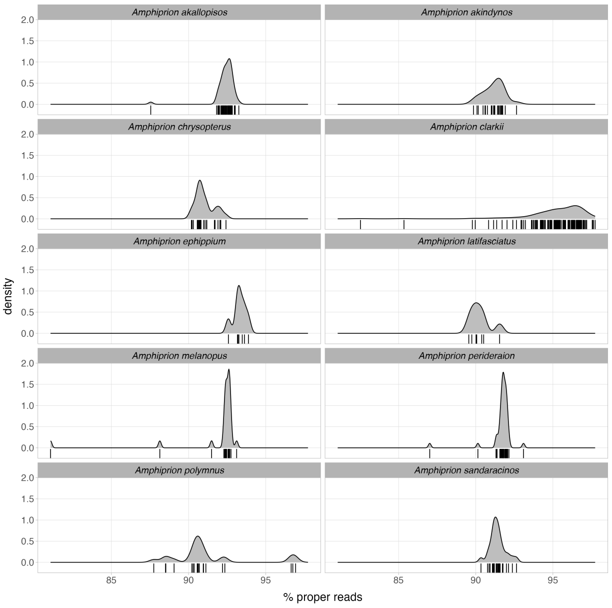


**b.**

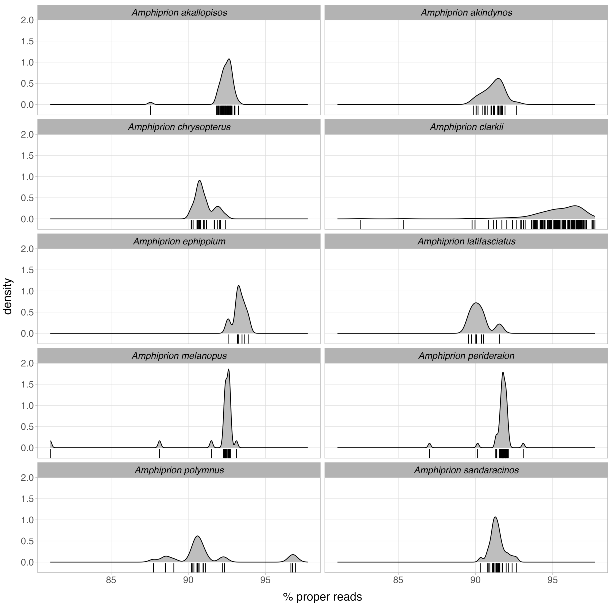


**Fig. S1**: Genomics summary per species. **a.** Density distribution of percentage of mapped proper reads per individual grouped by species. **b.** Density distribution of average sequencing depth per individual grouped by species.

**a.**

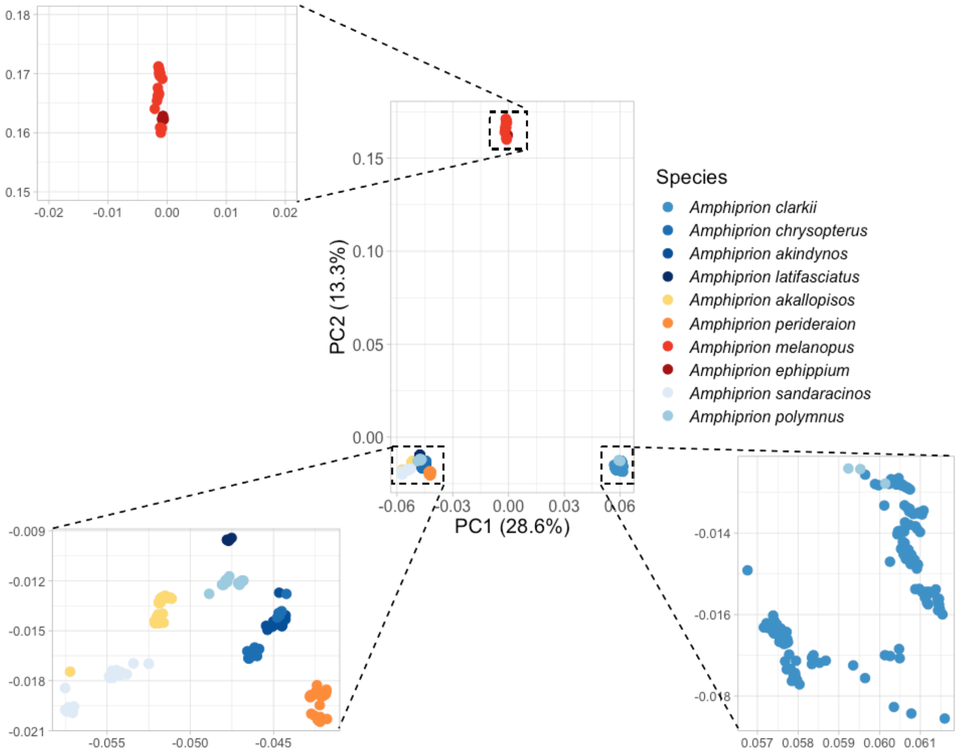


**b.**

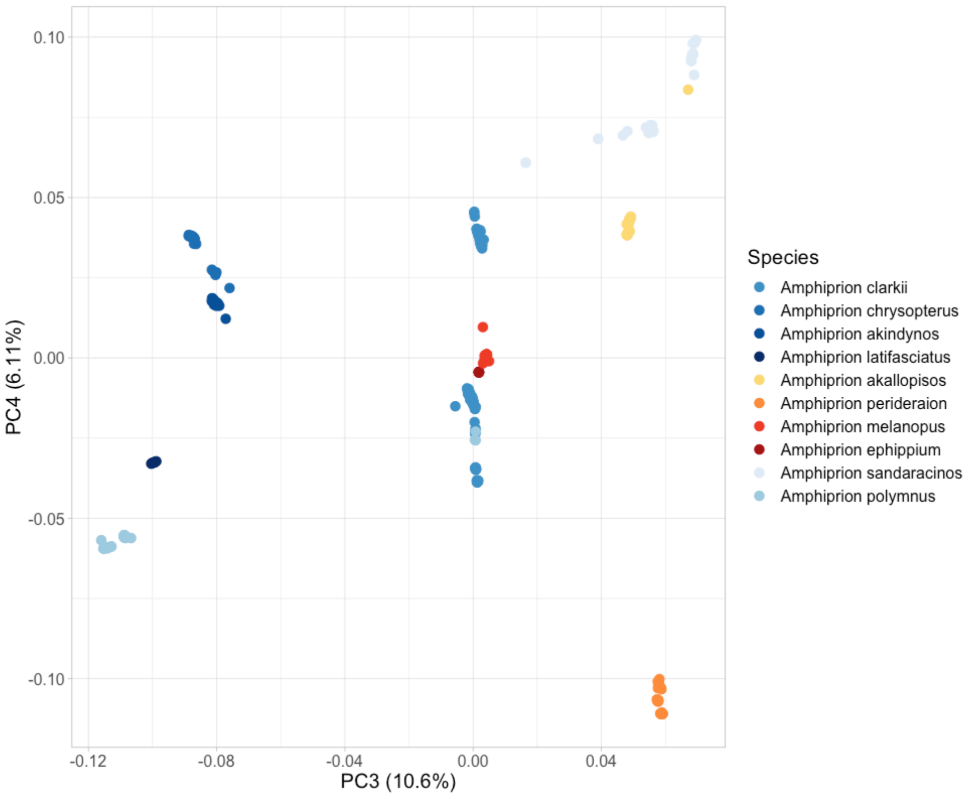


**Fig. S2**: PCA on all-species SNP dataset. **a.** PC1 and PC2 from Principal Component Analysis (PCA) on all-species SNP dataset. PC1 explains 28.6% and PC2 13.3% of the genetic variance across samples. Individuals are colored by species, shown in legend. **b.** PC3 and PC4 from Principal Component Analysis (PCA) on all-species SNP dataset. PC3 explains 10.6% and PC4 6.11% of the genetic variance across samples. Individuals are colored by species, shown in legend.

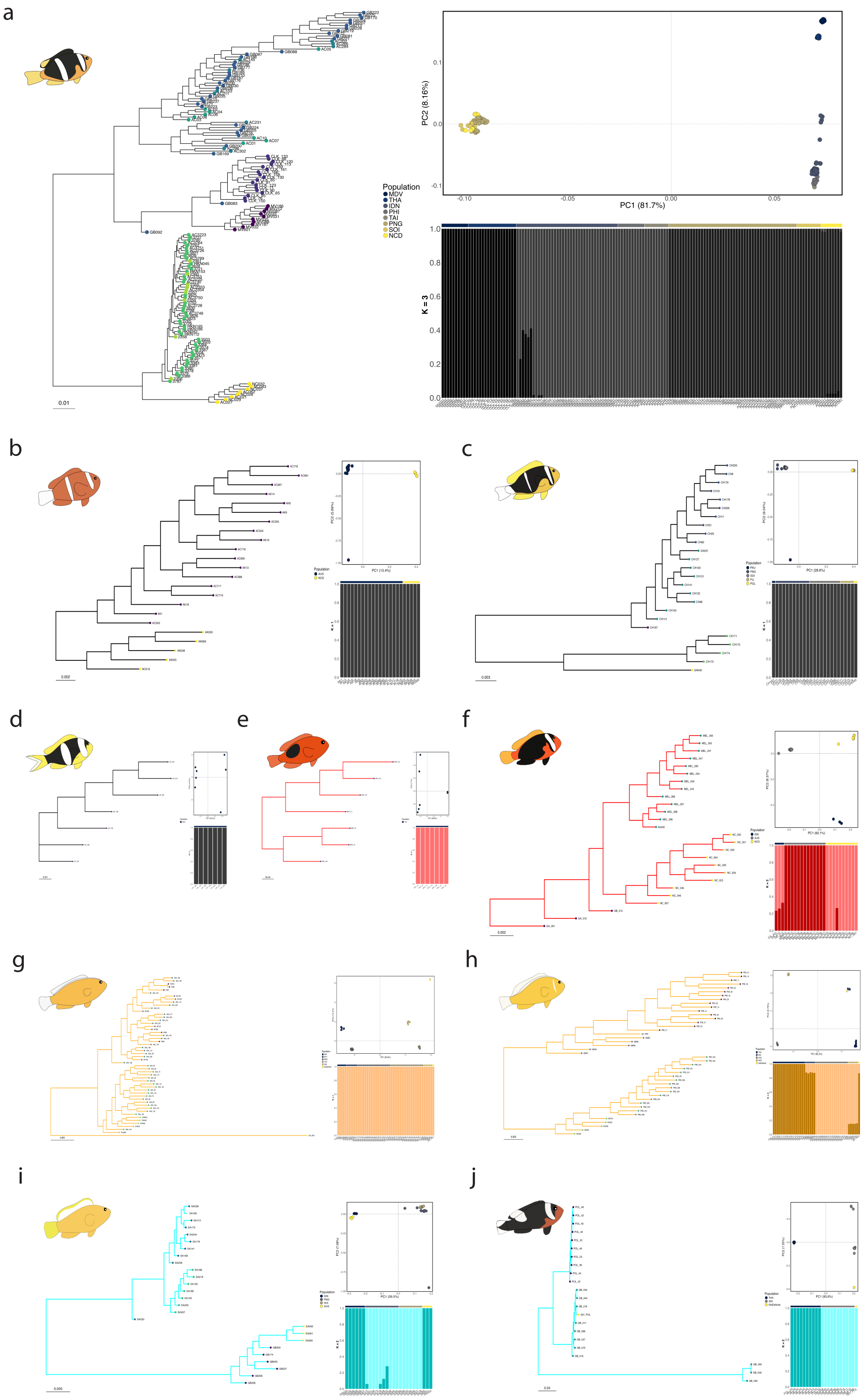


**Fig. S3**: Population structure of the 10 clownfish species. Color codes are according to their host regime (black for generalists, red for EQ specialists, orange for RM specialists and blue for SD specialists). Top-panel: Phylogenetic reconstruction with tips colored by population. Color code per population is shown at the top of the plot, Mid-panel: First and second principal components of the PCA. Proportion of variance explain for each axis is indicated at the axis label between parenthesis. Colored points are according to the population.; and Bottom-panel: ADMIXTURE plot showing the best K, i.e. the lowest cross-validation error. Colored bars at the top of each admixture plot indicate different populations with population names indicated by their abbreviation above the colored bar. Bar heights show the proportion of each individual's genome assigned to the color-coded genetic clusters. a) *A. clarkii*, b) *A. akindynos*, c) *A. chrysopterus*, d) *A. latifasciatus*, e) *A. ephippium*, f) *A. melanopus*, g) *A. akallopisos*, h) *A. perideraion*, i) *A. sandaracinos*, and j) *A. polymnus*.

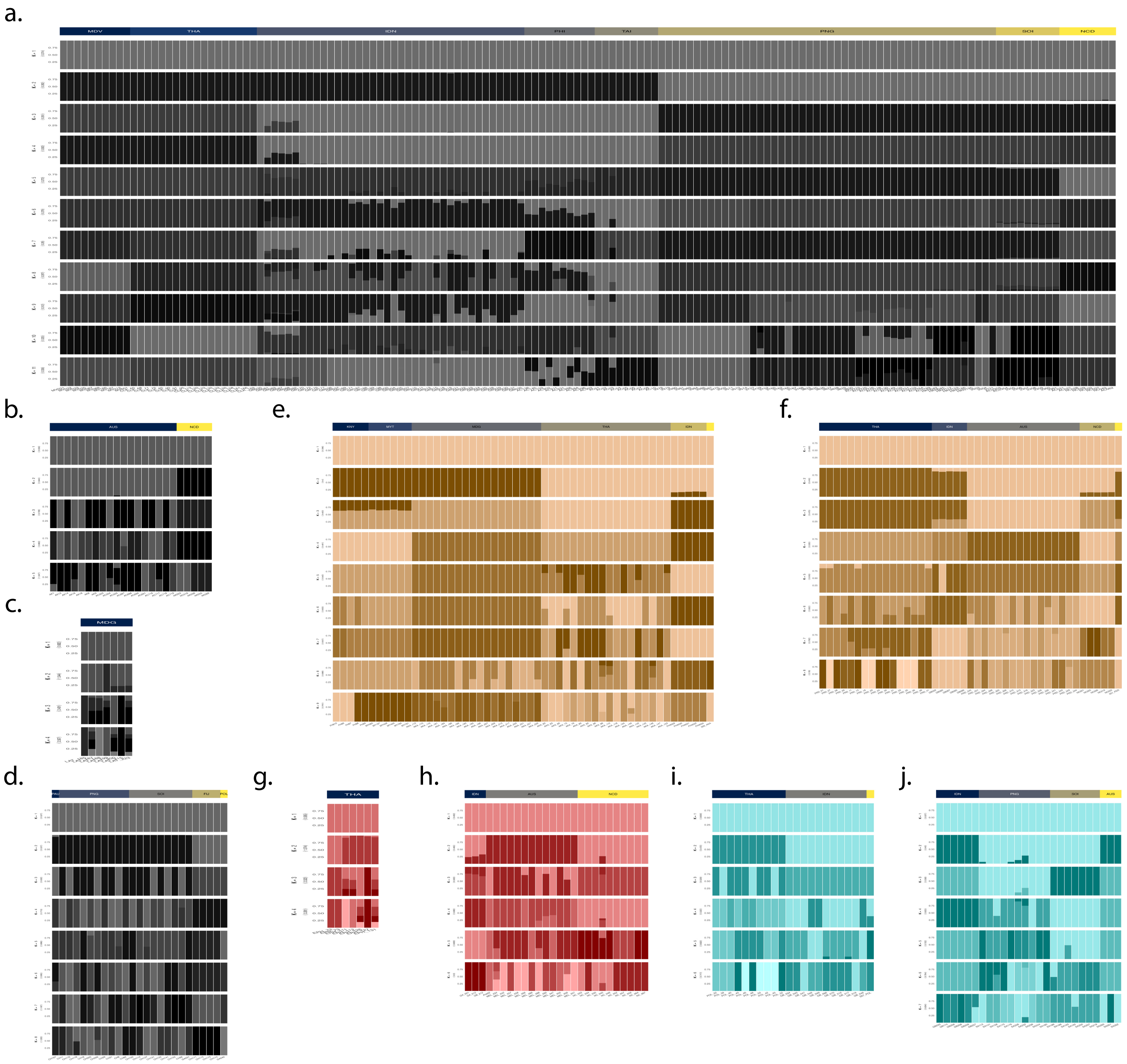


**Fig. S4**: Admixture plots of the 10 clownfish species. Color bars are colored according to their host regime (black for generalists, red for EQ specialists, orange for RM specialists and blue for SD specialists). For each plot, the different shades of the main color hue represent the distinct genetic clusters identified by ADMIXTURE when it was run with a specific number of clusters (K), indicated on the left of the plot. Cross validation error of each K is indicated within parenthesis below the number of clusters. Colored bars at the top of each admixture plot indicate different populations with population names indicated by their abbreviation inside the colored bar. Bar heights show the proportion of each individual's genome assigned to the color-coded genetic clusters. a) *A. clarkii*, b) *A. akindynos*, c) *A. latifasciatus*, d) *A. chrysopterus*, e) *A. akallopisos*, f) *A. perideraion*, g) *A. ephippium*, h) *A. melanopus*, i) *A. polymnus*, and j) *A. sandarancinos*.

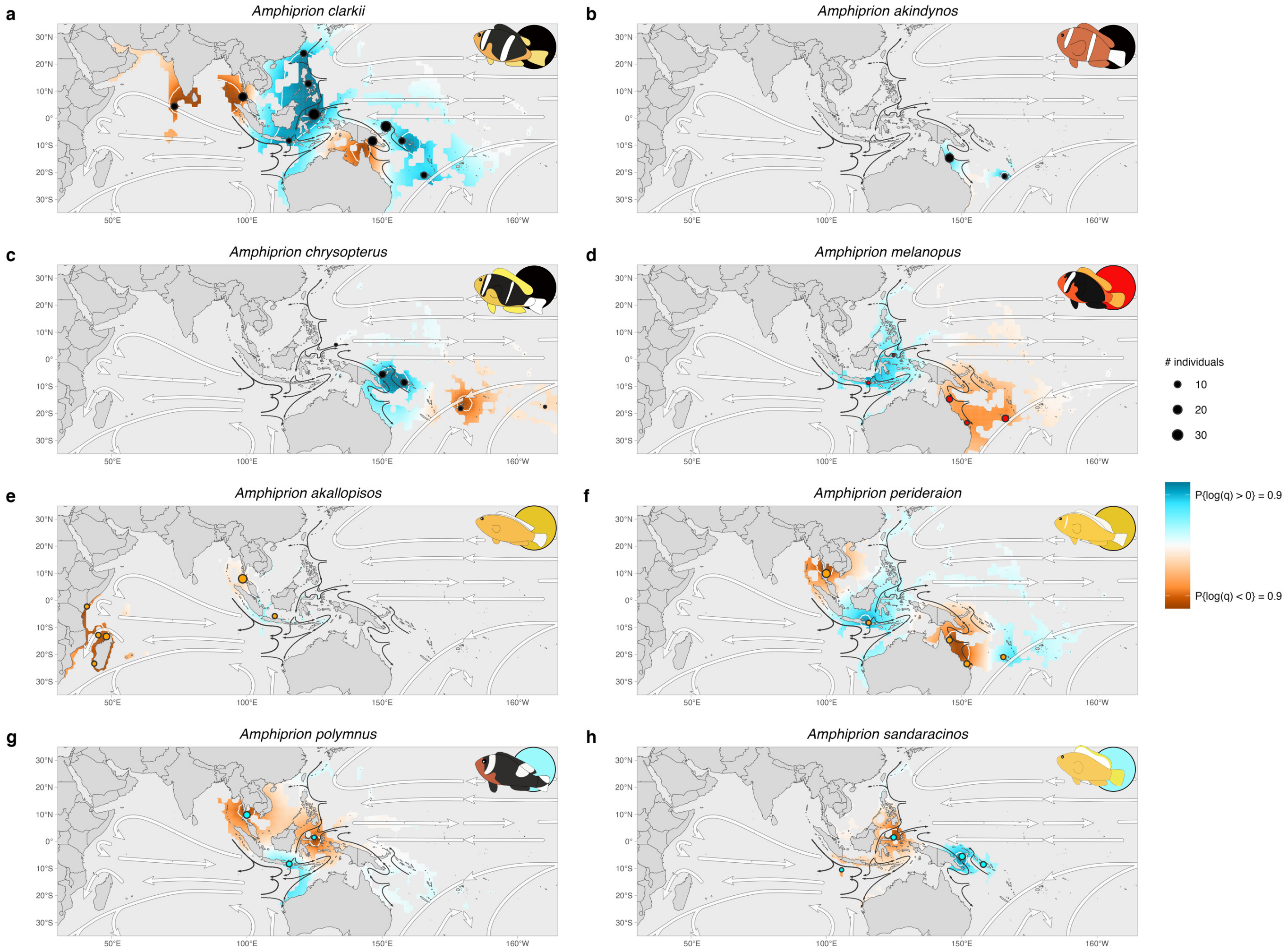


**Fig. S5**: Posterior probabilities of genetic diversity being higher or lower than the expected under the Isolation-by-distance model obtained from eems [**?** ]. Maps show the extent of the clownfish clade distribution (from Eastern Africa to the East Pacific and from Temperate Australia to Japan). Filled circles indicate the sampling locations with the size being the number of individuals and the color denoting the host category (black for generalists, orange for RM specialists, red for EQ specialists and blue for SD specialists). Genetic diversity surfaces are masked by each species distribution obtained from SDMs. White filled arrows denote global oceanic currents, while dark arrows indicate sea currents of the Coral Triangle. Values indicate the posterior probability of a location having higher (blue) or lower (red) genetic differentiation than expected under Isolation-by-distance.

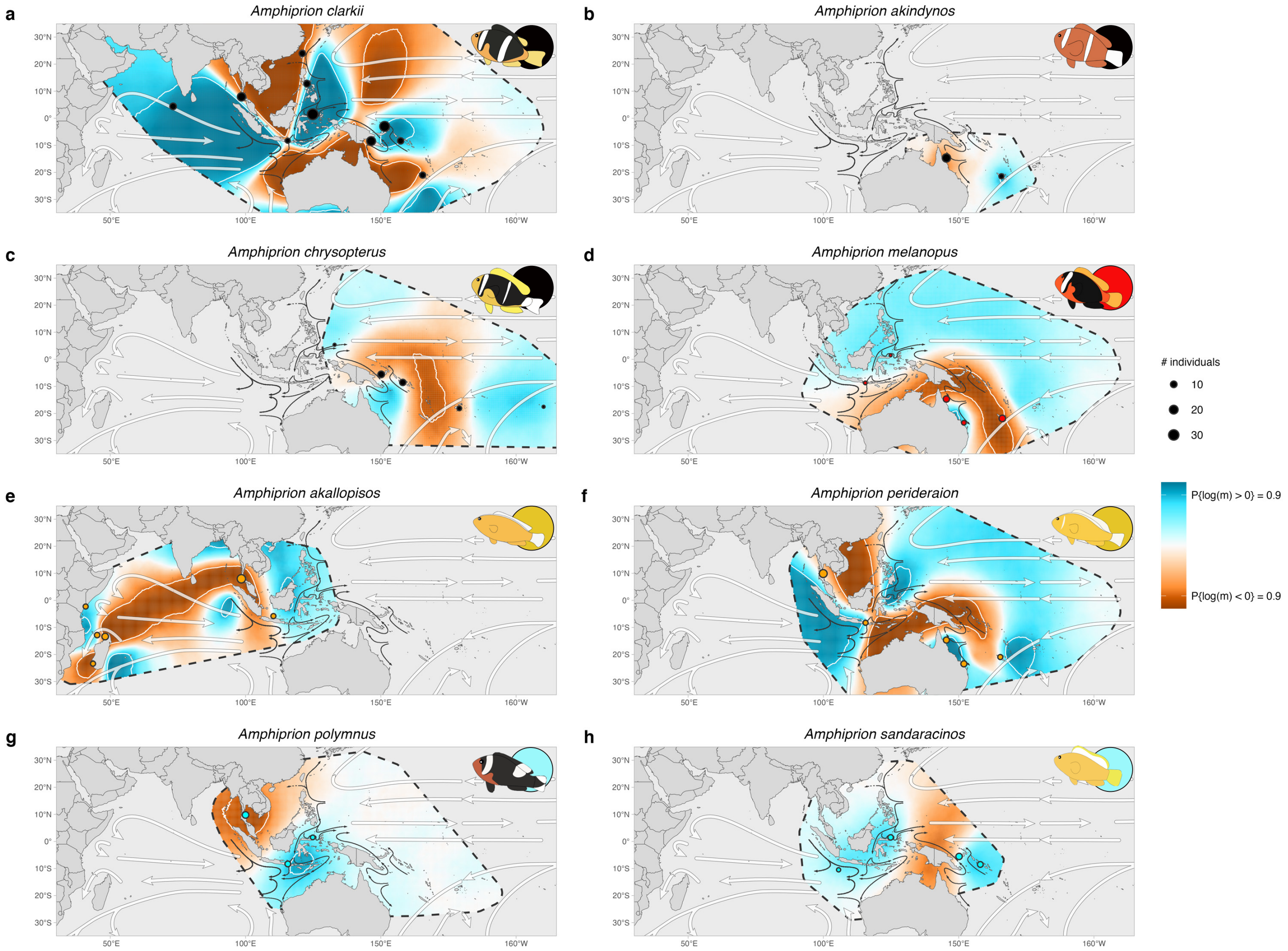


**Fig. S6**: Posterior probabilities of effective migration rate being higher or lower than the expected under Isolation-by-distance (IBD) obtained from eems [**?** ]. Maps show the extent of the clownfish clade distribution (from Eastern Africa to the East Pacific and from Temperate Australia to Japan). Filled circles indicate the sampling locations with the size being the number of individuals and the color denoting the host category (black for generalists, orange for RM specialists, red for EQ specialists and blue for SD specialists). White-filled arrows denote global oceanic currents, while dark arrows indicate sea currents of the Coral Triangle. Values indicate the posterior probability of a location having higher (blue) or lower (red) migration than expected under Isolation-by-distance (IBD).

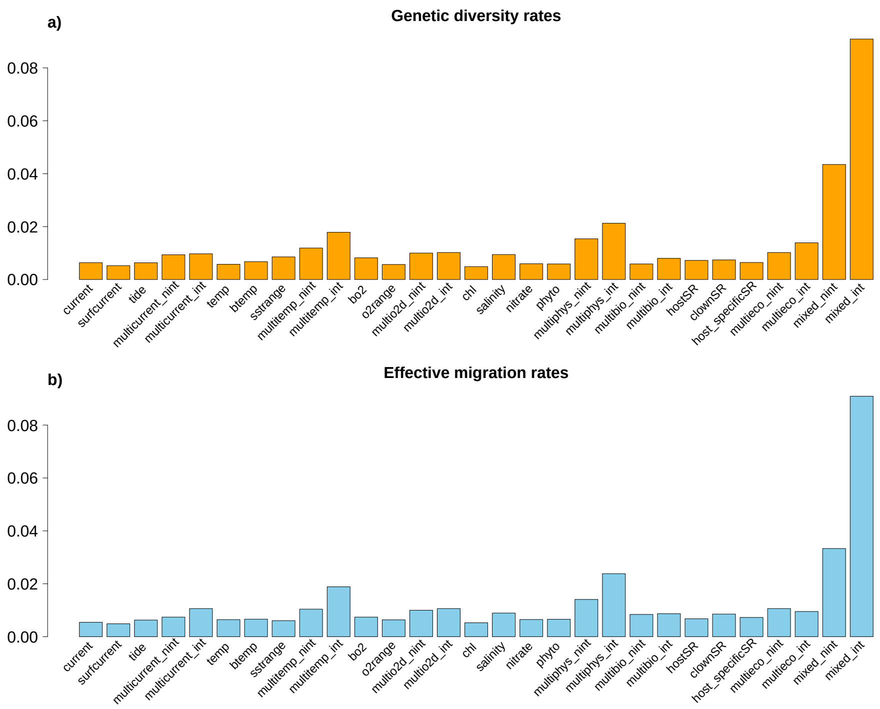


**Fig. S7**: Barplot indicating the ranking of multiple spatial auto-regressive models on (a) genetic diversity rates and (b) effective migration rates. The ranking for each model is computed as the inverse of the sum of its rankings across multiple species, where a higher bar represents a better overall ranking. This approach allows for a comparative assessment of model performance across different biological metrics, with the highest-ranked model being the one with the lowest aggregate rank across species.

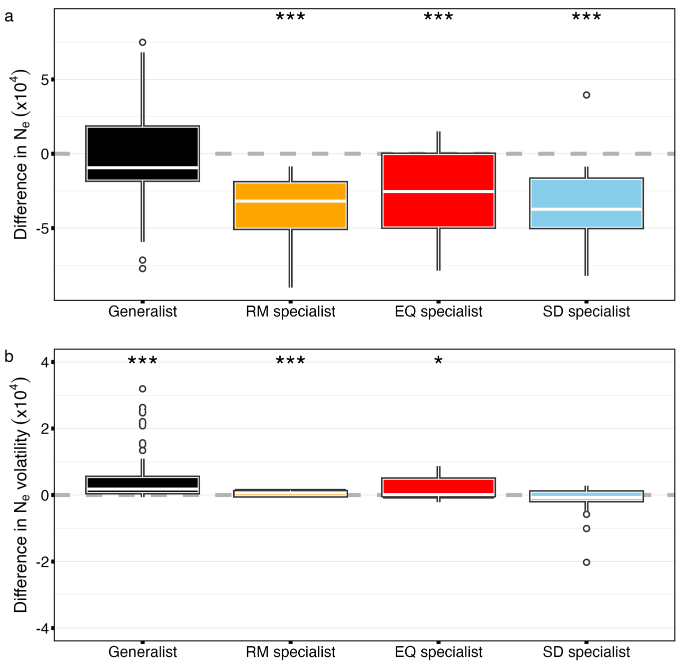


**Fig. S8**: Comparisons of cumulative differences over time among the distinct host specialization groups. a) Historical $N_e$. b) $N_e$ volatility. Grey dashed line denotes no cumulative differences over time. White line represents the median (Q2), while colored boxes represent the first to third quartile range (IQR). Whiskers show the upper and lower bounds. Legend at the top of each box plot indicates the statistical significance of the Wilcoxon test comparing each group's distribution to zero.

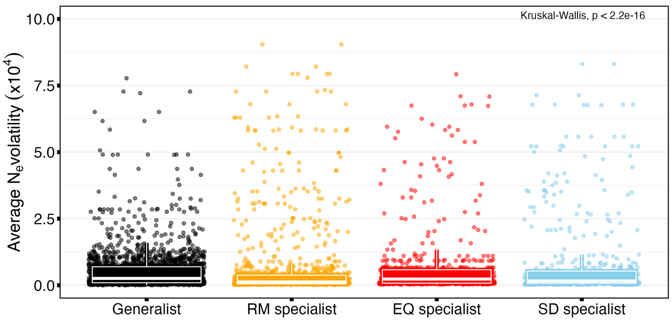


**Fig. S9**: Comparison of $N_e$ volatility values across host categories. $N_e$ volatility is calculated as the average true range (ATR) over 10,000-year intervals.

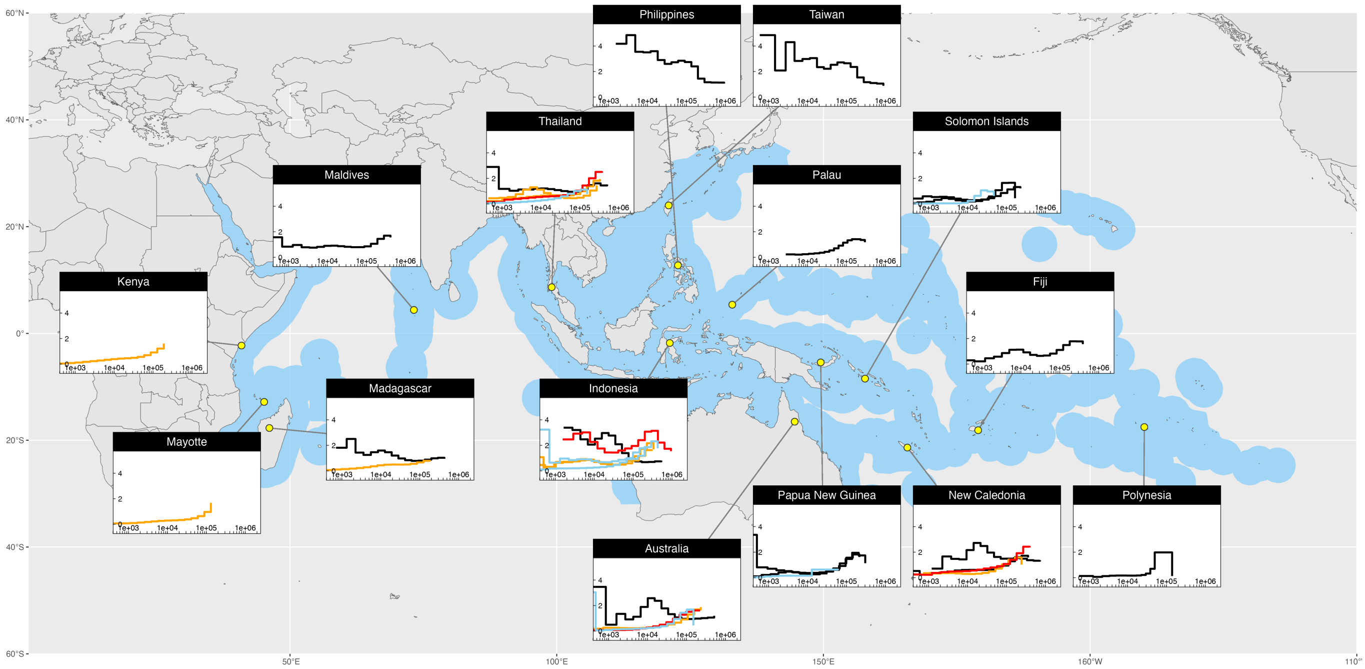


**Fig. S10**: Distribution map of clownfishes shown in light blue polygons over the Indo-Pacific with species MSMC reconstructions of each population (yellow filled-circle). Names of the country where the population was sampled is indicated at the top of each msmc plot. $N_e$ reconstruction lines are colored accordingly to the host specialization (black for Generalists, orange for *Radianthus* specialists, red for *Entacmaea* specialists, and blue for *Stichodactyla* specialists.). $N_e$ values are in the scale of $10^4$ and time axis shows a time frame from $10^3$ to $10^6$ years ago considering a mutation rate ($\mu$) of $4e^{-8}$ and a generation time of 5 years.

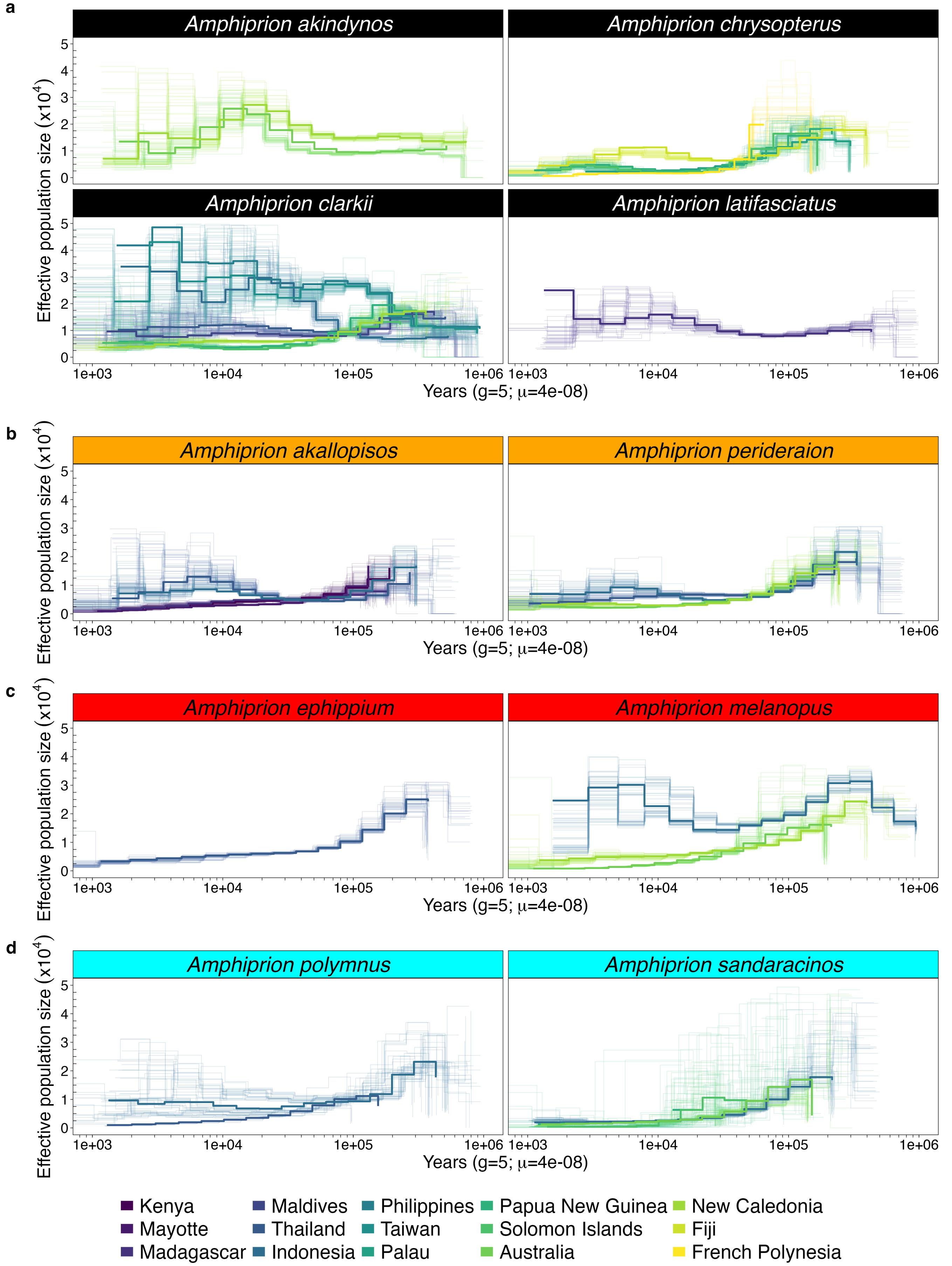


**Fig. S11**: Demographic reconstructions of all populations of each species. Thick colored line shows the average trajectory over ten msmc runs using 3 random individuals each time from each population. Light colored lines represent 50 bootstraps on each population using msmc-tools function 'multihetsep_bootstrap.py'. a) Generalists; b) *Radianthus magnifica* specialists (RM specialists); c) *Entacmaea quadricolor* specialists (EQ specialists); and d) *Stichodactyla* specialists (SD specialists).

### Supplementary Tables

**Tab. S1**: Species used in this study, number of individuals and populations and their mutualistic behavior (RM specialist: *Radianthus magnifica*; EQ specialist: *Entacmaea quadricolor*; SD specialist: *Stichodactyla* genus, including *S. mertensii*, *S. haddoni* and *S. magnifica*, and Generalist).

| Species | Num. individuals | Num. populations | Host classification |
|---|---|---|---|
| *Amphirion akallopisos* | 53 | 5 | RM specialist |
| *A. perideraion* | 43 | 4 | RM specialist |
| *A. melanopus* | 26 | 3 | EQ specialist |
| *A. ephippium* | 7 | 1 | EQ specialist |
| *A. sandaracinos* | 26 | 4 | SD specialist |
| *A. polymnus* | 22 | 2 | SD specialist |
| *A. akindynos* | 23 | 2 | Generalist |
| *A. chrysopterus* | 25 | 5 | Generalist |
| *A. clarkii* | 150 | 8 | Generalist |
| *A. latifasciatus* | 7 | 1 | Generalist |
| **Total** | 382 | | |

**Tab. S2**: Best K from admixture analysis along with number of defined populations per species, the geographical range computed from the species distribution models (SDMs), and the host category.

| species | best K | num. populations | geographic range ($km^2$) | host category |
|---|---|---|---|---|
| *Amphiprion akindynos* | 1 | 2 | 384992.345 | Generalist |
| *A. chrysopterus* | 1 | 5 | 550729.112 | Generalist |
| *A. clarkii* | 3 | 8 | 2175714.617 | Generalist |
| *A. latifasciatus* | 1 | 1 | 80968.679 | Generalist |
| *A. akallopisos* | 1 | 6 | 566945.887 | RM specialist |
| *A. perideraion* | 2 | 5 | 1533906.680 | RM specialist |
| *A. ephippium* | 1 | 1 | 232991.183 | EQ specialist |
| *A. melanopus* | 2 | 3 | 917404.851 | EQ specialist |
| *A. polymnus* | 2 | 3 | 1319327.140 | SD specialist |
| *A. sandaracinos* | 2 | 5 | 727854.832 | SD specialist |

**Tab. S3**: Spatial model specifications

| model | dependent variable | explanatory variables |
|---|---|---|
| current | $\log(q \mid m)$ | Current Velocity |
| surfcurrent | $\log(q \mid m)$ | Surface_current |
| tide | $\log(q \mid m)$ | Tide AVG |
| multicurrent_nint | $\log(q \mid m)$ | Current Velocity + Surface_current + Tide.AVG |
| multicurrent_int | $\log(q \mid m)$ | Current Velocity * Surface_current * Tide.AVG |
| temp | $\log(q \mid m)$ | Temperature |
| btemp | $\log(q \mid m)$ | Bottom temperature |
| sstrange | $\log(q \mid m)$ | SST range |
| multitemp_nint | $\log(q \mid m)$ | Temperature + Bottom temperature + SST.range |
| multitemp_int | $\log(q \mid m)$ | Temperature * Bottom temperature * SST.range |
| bo2 | $\log(q \mid m)$ | Bottom dO2 |
| o2range | $\log(q \mid m)$ | dO2 Range |
| multio2d_nint | $\log(q \mid m)$ | Bottom dO2 + dO2.Range |
| multio2d_int | $\log(q \mid m)$ | Bottom dO2 * dO2.Range |
| chl | $\log(q \mid m)$ | Chl |
| salinity | $\log(q \mid m)$ | Salinity |
| nitrate | $\log(q \mid m)$ | Nitrate |
| phyto | $\log(q \mid m)$ | Phytoplankton |
| multiphys_nint | $\log(q \mid m)$ | Current Velocity + Temperature + Salinity |
| multiphys_int | $\log(q \mid m)$ | Current Velocity * Temperature * Salinity |
| multibio_nint | $\log(q \mid m)$ | Chl + Nitrate + Phytoplankton |
| multibio_int | $\log(q \mid m)$ | Chl * Nitrate * Phytoplankton |
| hostSR | $\log(q \mid m)$ | hostRichness |
| clownSR | $\log(q \mid m)$ | clownfishRichness |
| host_specificSR | $\log(q \mid m)$ | hostSpecRich |
| multieco_nint | $\log(q \mid m)$ | hostRichness + clownfishRichness + hostSpecRich |
| multieco_int | $\log(q \mid m)$ | hostRichness * clownfishRichness * hostSpecRich |
| mixed_nint | $\log(q \mid m)$ | Bottom dO2 + SST.range + Current.Velocity + Tide.AVG + Phytoplankton + hostRichness + clownfishRichness |
| mixed_int | $\log(q \mid m)$ | Bottom dO2 * SST.range * Current.Velocity * Tide.AVG * Phytoplankton * hostRichness * clownfishRichness |

**Tab. S4**: AICc of species spatial auto-regressive models on genetic diversity. Number of parameters (K) of each model are specified in the second column. Three best models based on the lowest AICc are indicated in bold with their rank indicated between parenthesis.

| Model | K | A. akindynos | A. chrysopterus | A. clarkii | A. akallopisos | A. perideraion | A. melanopus | A. polymnus | A. sandaracinos |
|---|---|---|---|---|---|---|---|---|---|
| current | 4 | -6202.915 | -3662.758 | -3564.486 | -3288.253 | -5123.984 | -4486.163 | -4812.212 | -3039.973 |
| surfcurrent | 4 | -6234.956 | -3599.253 | -3564.171 | -3154.704 | -5113.202 | -4505.071 | -4806.33 | -3025.28 |
| tide | 4 | -6226.923 | -3570.718 | -3568.282 | -3198.714 | -5113.04 | -4558.965 | -4851.797 | -3059.274 |
| multicurrent_nint | 6 | -6302.693 | -3684.399 | -3564.679 | -3303.891 | -5120.674 | -4574.781 | -4856.776 | -3069.737 |
| multicurrent_int | 10 | -6325.886 | -3686.355 | -3561.279 | -3349.554 | -5123.136 | -4593.749 | -4889.553 | -3066.626 |
| temp | 4 | -6183.982 | -3598.61 | -3564.231 | -3200.555 | -5144.752 | -4503.026 | -4838.513 | -3023.296 |
| btemp | 4 | -6185.75 | -3619.307 | -3564.237 | -3222.524 | -5195.244 | -4622.333 | -4825.815 | -3032.056 |
| sstrange | 4 | -6185.849 | -3570.719 | -3605.336 | -3591.078 | -5147.745 | -4841.486 | -4935.045 | -3041.168 |
| multitemp_nint | 6 | -6184.784 | -3666.273 | -3605.533 | -3625.813 | -5212.844 | -4841.578 | -4974.254 | -3058.396 |
| multitemp_int | 10 | -6231.842 | -3687.324 | -3624.695 | **-3694.664(3)** | -5229.17 | -4864.223 | **-4989.501(3)** | -3059.465 |
| bo2 | 4 | -6203.605 | -3572.472 | -3631.098 | -3234.758 | -5222.079 | -5001.095 | -4807.346 | -3043.682 |
| o2range | 4 | -6290.157 | -3622.928 | -3565.149 | -3206.346 | -5113.216 | -4488.016 | -4807.548 | -3021.963 |
| multio2d_nint | 5 | -6289.666 | -3631.018 | -3629.092 | -3263.845 | -5223.296 | -5010.27 | -4806.4 | -3042.514 |
| multio2d_int | 6 | -6306.364 | -3629.229 | -3637.405 | -3278.9 | -5222.328 | **-5017.753(3)** | -4804.657 | -3040.515 |
| chl | 4 | -6183.848 | -3573.602 | -3571.651 | -3151.59 | -5113.813 | -4486.154 | -4813.41 | -3022.339 |
| salinity | 4 | -6447.101 | -3570.885 | **-3694.643(2)** | -3519.397 | -5115.156 | -4854.72 | -4816.582 | -3116.903 |
| nitrate | 4 | -6195.996 | -3587.854 | -3564.925 | -3152.185 | -5161.815 | -4554.712 | -4828.294 | -3024.129 |
| phyto | 4 | -6191.299 | -3589.922 | -3566.59 | -3168.033 | -5141.181 | -4515.76 | -4834.723 | -3023.688 |
| multiphys_nint | 6 | -6500.544 | -3667.39 | **-3692.953(3)** | -3552.457 | -5154.31 | -4853.382 | -4837.118 | **-3152.369(2)** |
| multiphys_int | 10 | -6514.666 | **-3694.543(3)** | **-3782.294(1)** | -3579.746 | **-5318.639(2)** | -4862.439 | -4834.239 | **-3152.089(3)** |
| multibio_nint | 6 | -6192.705 | -3587.574 | -3567.718 | -3179.996 | -5158.014 | -4552.847 | -4831.912 | -3021.696 |
| multibio_int | 10 | -6306.099 | -3601.137 | -3584.184 | -3205.293 | -5164.296 | -4642.929 | -4838.184 | -3022.254 |
| hostSR | 4 | -6264.32 | -3590.008 | -3581.378 | -3301.317 | -5123.341 | -4493.718 | -4854.805 | -3032.903 |
| clownSR | 4 | -6536.638 | -3616.956 | -3575.52 | -3404.692 | -5114.852 | -4523.902 | -4819.934 | -3032.089 |
| host_specificSR | 4 | -6275.805 | -3592.99 | -3581.378 | -3154.605 | -5119.173 | -4513.512 | -4846.357 | -3026.694 |
| multieco_nint | 6 | -6550.271 | -3629.138 | -3579.482 | -3591.743 | -5119.678 | -4560.942 | -4859.501 | -3054.2 |
| multieco_int | 10 | **-6674.098(2)** | -3661.705 | -3621.489 | -3638.897 | -5135.357 | -4572.894 | -4872.756 | -3101.055 |
| mixed_nint | 10 | **-6652.014(3)** | **-3710.708(2)** | -3678.047 | **-3834.389(2)** | **-5250.199(3)** | **-5115.779(2)** | **-5045.714(2)** | -3137.381 |
| mixed_int | 130 | **-7348.41(1)** | **-3850.82(1)** | -3685.674 | **-4088.833(1)** | **-5473.507(1)** | **-5292.367(1)** | **-5222.384(1)** | **-3189.784(1)** |

**Tab. S5**: AICc of species spatial auto-regressive models on effective migration rates. Number of parameters (K) per model are specified in the second column.Three best models based on the lowest AICc are indicated in bold with their rank indicated between parenthesis.

| Model | K | A. akindynos | A. chrysopterus | A. clarkii | A. akallopisos | A. perideraion | A. melanopus | A. polymnus | A. sandaracinos |
|---|---|---|---|---|---|---|---|---|---|
| current | 4 | 340.52 | 673.798 | 2182.525 | 1930.811 | 1683.695 | 1208.978 | 1159.288 | 509.836 |
| surfcurrent | 4 | 411.377 | 673.845 | 2182.606 | 1931.228 | 1693.284 | 1209.882 | 1161.158 | 511.883 |
| tide | 4 | 360.245 | 591.098 | 2158.761 | 1937.86 | 1682.705 | 1209.37 | 1145.967 | 511.099 |
| multicurrent_nint | 6 | 307.284 | 586.108 | 2154.232 | 1928.479 | 1668.833 | 1210.24 | 1143.563 | 512.031 |
| multicurrent_int | 10 | 283.689 | **540.519(3)** | **2115.382(3)** | 1930.225 | 1652.897 | 1174.712 | 1140.132 | 499.315 |
| temp | 4 | 257.138 | 668.502 | 2183.64 | 1932.039 | 1681.966 | 1205.777 | 1137.68 | 487.18 |
| btemp | 4 | 244.889 | 666.35 | 2184.126 | 1938.688 | 1671.992 | 1209.039 | 1121.737 | 486.227 |
| sstrange | 4 | 412.653 | 658.167 | 2158.732 | 1926.024 | 1672.072 | 1210.533 | 1161.149 | 496.285 |
| multitemp_nint | 6 | 207.292 | 652.138 | 2154.845 | 1918.007 | 1660.202 | 1209.469 | 1125.066 | 458.176 |
| multitemp_int | 10 | 186.569 | 639.626 | 2123.179 | 1888.201 | 1661.534 | 1159.68 | 1064.04 | **431.858(3)** |
| bo2 | 4 | 326.486 | 579.934 | 2157.812 | 1935.802 | 1691.802 | 1192.95 | 1092.937 | 512.378 |
| o2range | 4 | 237.851 | 671.61 | 2179.035 | 1917.378 | 1696.523 | 1208.951 | 1153.311 | 510.969 |
| multio2d_nint | 5 | **165.285(3)** | 567.365 | 2148.306 | 1919.235 | 1693.833 | 1192.509 | 1087.868 | 512.922 |
| multio2d_int | 6 | 166.226 | 568.495 | 2150.327 | 1921.218 | 1677.887 | 1185.875 | 1088.403 | 512.443 |
| chl | 4 | 373.675 | 673.979 | 2181.064 | 1938.469 | 1687.435 | 1207.97 | 1150.585 | 507.38 |
| salinity | 4 | 408.603 | 640.691 | 2180.584 | 1937.584 | 1671.736 | 1160.717 | 1041.631 | 438.295 |
| nitrate | 4 | 243.833 | 654.422 | 2184.571 | 1938.123 | 1677.623 | 1205.925 | 1136.035 | 494.239 |
| phyto | 4 | 269.348 | 668.99 | 2182.07 | 1932.605 | 1673.544 | 1210.288 | 1111.92 | 485.774 |
| multiphys_nint | 6 | 253.659 | 626.714 | 2178.961 | 1912.753 | 1657.533 | **1120.241(2)** | 1044.869 | 441.496 |
| multiphys_int | 10 | 224.899 | 588.478 | 2175.266 | **1718.485(2)** | **1596.969(2)** | **1123.095(3)** | **984.116(2)** | **369.25(1)** |
| multibio_nint | 6 | 246.096 | 623.77 | 2181.56 | 1924.448 | 1674.044 | 1204.699 | 1115.541 | 489.641 |
| multibio_int | 10 | 248.193 | 619.355 | 2180.692 | 1910.66 | 1675.932 | 1209.913 | 1112.539 | 482.761 |
| hostSR | 4 | 310.217 | 654.047 | 2172.59 | 1923.744 | 1685.393 | 1211.08 | 1067.802 | 501.327 |
| clownSR | 4 | 362.369 | 650.269 | 2161.129 | 1909.115 | 1660.655 | 1208.43 | 1124.869 | 487.598 |
| host_specificSR | 4 | 332.486 | 648.721 | 2172.59 | 1925.644 | 1661.236 | 1210.696 | 1132.247 | 485.882 |
| multieco_nint | 6 | 307.095 | 648.663 | 2163.149 | 1837.949 | 1637.613 | 1209.976 | 1068.545 | 478.016 |
| multieco_int | 10 | 313.333 | 655.609 | 2168.454 | **1820.063(3)** | **1617.58(3)** | 1216.949 | 1058.594 | 481.623 |
| mixed_nint | 10 | **137.138(2)** | **370.875(2)** | **2103.757(2)** | 1874.869 | 1626.21 | 1187.873 | **950.753(1)** | 451.873 |
| mixed_int | 130 | **-168.014(1)** | **283.342(1)** | **1884.342(1)** | **1694.958(1)** | **1490.122(1)** | **1040.961(1)** | **998.813(3)** | **377.154(2)** |

**Tab. S6**: Model selection table of Lagrange Multiplier (LM) Spatial Auto-Regressive (SAR) models of *A. akallopisos* on inferred deviations from genetic diversity expectations under Isolation-by-distance

| Model | n | K | AIC | AICc | Delta.AICc | AICcWt | Cum.Wt | LogLik |
|---|---|---|---|---|---|---|---|---|
| mixed_int | 598 | 130.00 | -4161.77 | -4088.83 | 0.00 | 1.00 | 1.00 | 2209.88 |
| mixed_nint | 598 | 10.00 | -3834.76 | -3834.39 | 254.44 | 0.00 | 1.00 | 1926.38 |
| multitemp_int | 598 | 10.00 | -3695.04 | -3694.66 | 394.17 | 0.00 | 1.00 | 1856.52 |
| multieco_int | 598 | 10.00 | -3639.27 | -3638.90 | 449.94 | 0.00 | 1.00 | 1828.64 |
| multitemp_nint | 598 | 6.00 | -3625.95 | -3625.81 | 463.02 | 0.00 | 1.00 | 1817.98 |
| multieco_nint | 598 | 6.00 | -3591.89 | -3591.74 | 497.09 | 0.00 | 1.00 | 1800.94 |
| sstrange | 598 | 4.00 | -3591.15 | -3591.08 | 497.75 | 0.00 | 1.00 | 1798.57 |
| multiphys_int | 598 | 10.00 | -3580.12 | -3579.75 | 509.09 | 0.00 | 1.00 | 1799.06 |
| multiphys_nint | 598 | 6.00 | -3552.60 | -3552.46 | 536.38 | 0.00 | 1.00 | 1781.30 |
| salinity | 598 | 4.00 | -3519.46 | -3519.40 | 569.44 | 0.00 | 1.00 | 1762.73 |
| clownSR | 598 | 4.00 | -3404.76 | -3404.69 | 684.14 | 0.00 | 1.00 | 1705.38 |
| multicurrent_int | 598 | 10.00 | -3349.93 | -3349.55 | 739.28 | 0.00 | 1.00 | 1683.96 |
| multicurrent_nint | 598 | 6.00 | -3304.03 | -3303.89 | 784.94 | 0.00 | 1.00 | 1657.02 |
| hostSR | 598 | 4.00 | -3301.38 | -3301.32 | 787.52 | 0.00 | 1.00 | 1653.69 |
| current | 598 | 4.00 | -3288.32 | -3288.25 | 800.58 | 0.00 | 1.00 | 1647.16 |
| multio2d_int | 598 | 6.00 | -3279.04 | -3278.90 | 809.93 | 0.00 | 1.00 | 1644.52 |
| multio2d_nint | 598 | 5.00 | -3263.95 | -3263.84 | 824.99 | 0.00 | 1.00 | 1635.97 |
| bo2 | 598 | 4.00 | -3234.83 | -3234.76 | 854.08 | 0.00 | 1.00 | 1620.41 |
| btemp | 598 | 4.00 | -3222.59 | -3222.52 | 866.31 | 0.00 | 1.00 | 1614.30 |
| o2range | 598 | 4.00 | -3206.41 | -3206.35 | 882.49 | 0.00 | 1.00 | 1606.21 |
| multibio_int | 598 | 10.00 | -3205.67 | -3205.29 | 883.54 | 0.00 | 1.00 | 1611.83 |
| temp | 598 | 4.00 | -3200.62 | -3200.55 | 888.28 | 0.00 | 1.00 | 1603.31 |
| tide | 598 | 4.00 | -3198.78 | -3198.71 | 890.12 | 0.00 | 1.00 | 1602.39 |
| multibio_nint | 598 | 6.00 | -3180.14 | -3180.00 | 908.84 | 0.00 | 1.00 | 1595.07 |
| phyto | 598 | 4.00 | -3168.10 | -3168.03 | 920.80 | 0.00 | 1.00 | 1587.05 |
| surfcurrent | 598 | 4.00 | -3154.77 | -3154.70 | 934.13 | 0.00 | 1.00 | 1580.39 |
| host_specificSR | 598 | 4.00 | -3154.67 | -3154.60 | 934.23 | 0.00 | 1.00 | 1580.34 |
| nitrate | 598 | 4.00 | -3152.25 | -3152.19 | 936.65 | 0.00 | 1.00 | 1579.13 |
| chl | 598 | 4.00 | -3151.66 | -3151.59 | 937.24 | 0.00 | 1.00 | 1578.83 |

**Tab. S7**: Model selection table of Lagrange Multiplier (LM) Spatial Auto-Regressive (SAR) models of *A. akindynos* on inferred deviations from genetic diversity expectations under Isolation-by-distance

| Model | n | K | AIC | AICc | Delta.AICc | AICcWt | Cum.Wt | LogLik |
|---|---|---|---|---|---|---|---|---|
| mixed_int | 1000 | 130.00 | -7387.60 | -7348.41 | 0.00 | 1.00 | 1.00 | 3822.80 |
| multieco_int | 1000 | 10.00 | -6674.32 | -6674.10 | 674.31 | 0.00 | 1.00 | 3346.16 |
| mixed_nint | 1000 | 10.00 | -6652.24 | -6652.01 | 696.40 | 0.00 | 1.00 | 3335.12 |
| multieco_nint | 1000 | 6.00 | -6550.36 | -6550.27 | 798.14 | 0.00 | 1.00 | 3280.18 |
| clownSR | 1000 | 4.00 | -6536.68 | -6536.64 | 811.77 | 0.00 | 1.00 | 3271.34 |
| multiphys_int | 1000 | 10.00 | -6514.89 | -6514.67 | 833.74 | 0.00 | 1.00 | 3266.44 |
| multiphys_nint | 1000 | 6.00 | -6500.63 | -6500.54 | 847.87 | 0.00 | 1.00 | 3255.31 |
| salinity | 1000 | 4.00 | -6447.14 | -6447.10 | 901.31 | 0.00 | 1.00 | 3226.57 |
| multicurrent_int | 1000 | 10.00 | -6326.11 | -6325.89 | 1022.52 | 0.00 | 1.00 | 3172.05 |
| multio2d_int | 1000 | 6.00 | -6306.45 | -6306.36 | 1042.05 | 0.00 | 1.00 | 3158.22 |
| multibio_int | 1000 | 10.00 | -6306.32 | -6306.10 | 1042.31 | 0.00 | 1.00 | 3162.16 |
| multicurrent_nint | 1000 | 6.00 | -6302.78 | -6302.69 | 1045.72 | 0.00 | 1.00 | 3156.39 |
| o2range | 1000 | 4.00 | -6290.20 | -6290.16 | 1058.25 | 0.00 | 1.00 | 3148.10 |
| multio2d_nint | 1000 | 5.00 | -6289.73 | -6289.67 | 1058.74 | 0.00 | 1.00 | 3148.86 |
| host_specificSR | 1000 | 4.00 | -6275.85 | -6275.81 | 1072.61 | 0.00 | 1.00 | 3140.92 |
| hostSR | 1000 | 4.00 | -6264.36 | -6264.32 | 1084.09 | 0.00 | 1.00 | 3135.18 |
| surfcurrent | 1000 | 4.00 | -6235.00 | -6234.96 | 1113.45 | 0.00 | 1.00 | 3120.50 |
| multitemp_int | 1000 | 10.00 | -6232.06 | -6231.84 | 1116.57 | 0.00 | 1.00 | 3125.03 |
| tide | 1000 | 4.00 | -6226.96 | -6226.92 | 1121.49 | 0.00 | 1.00 | 3116.48 |
| bo2 | 1000 | 4.00 | -6203.65 | -6203.61 | 1144.80 | 0.00 | 1.00 | 3104.82 |
| current | 1000 | 4.00 | -6202.96 | -6202.92 | 1145.50 | 0.00 | 1.00 | 3104.48 |
| nitrate | 1000 | 4.00 | -6196.04 | -6196.00 | 1152.41 | 0.00 | 1.00 | 3101.02 |
| multibio_nint | 1000 | 6.00 | -6192.79 | -6192.71 | 1155.71 | 0.00 | 1.00 | 3101.39 |
| phyto | 1000 | 4.00 | -6191.34 | -6191.30 | 1157.11 | 0.00 | 1.00 | 3098.67 |
| sstrange | 1000 | 4.00 | -6185.89 | -6185.85 | 1162.56 | 0.00 | 1.00 | 3095.94 |
| btemp | 1000 | 4.00 | -6185.79 | -6185.75 | 1162.66 | 0.00 | 1.00 | 3095.90 |
| multitemp_nint | 1000 | 6.00 | -6184.87 | -6184.78 | 1163.63 | 0.00 | 1.00 | 3097.43 |
| temp | 1000 | 4.00 | -6184.02 | -6183.98 | 1164.43 | 0.00 | 1.00 | 3095.01 |
| chl | 1000 | 4.00 | -6183.89 | -6183.85 | 1164.56 | 0.00 | 1.00 | 3094.94 |

**Tab. S8**: Model selection table of Lagrange Multiplier (LM) Spatial Auto-Regressive (SAR) models of *A. chrysopterus* on inferred deviations from genetic diversity expectations under Isolation-by-distance

| Model | n | K | AIC | AICc | Delta.AICc | AICcWt | Cum.Wt | LogLik |
|---|---|---|---|---|---|---|---|---|
| mixed_int | 1000 | 130.00 | -3890.01 | -3850.82 | 0.00 | 1.00 | 1.00 | 2074.01 |
| mixed_nint | 1000 | 10.00 | -3710.93 | -3710.71 | 140.11 | 0.00 | 1.00 | 1864.47 |
| multiphys_int | 1000 | 10.00 | -3694.77 | -3694.54 | 156.28 | 0.00 | 1.00 | 1856.38 |
| multitemp_int | 1000 | 10.00 | -3687.55 | -3687.32 | 163.50 | 0.00 | 1.00 | 1852.77 |
| multicurrent_int | 1000 | 10.00 | -3686.58 | -3686.35 | 164.47 | 0.00 | 1.00 | 1852.29 |
| multicurrent_nint | 1000 | 6.00 | -3684.48 | -3684.40 | 166.42 | 0.00 | 1.00 | 1847.24 |
| multiphys_nint | 1000 | 6.00 | -3667.47 | -3667.39 | 183.43 | 0.00 | 1.00 | 1838.74 |
| multitemp_nint | 1000 | 6.00 | -3666.36 | -3666.27 | 184.55 | 0.00 | 1.00 | 1838.18 |
| current | 1000 | 4.00 | -3662.80 | -3662.76 | 188.06 | 0.00 | 1.00 | 1834.40 |
| multieco_int | 1000 | 10.00 | -3661.93 | -3661.71 | 189.11 | 0.00 | 1.00 | 1839.96 |
| multio2d_nint | 1000 | 5.00 | -3631.08 | -3631.02 | 219.80 | 0.00 | 1.00 | 1819.54 |
| multio2d_int | 1000 | 6.00 | -3629.31 | -3629.23 | 221.59 | 0.00 | 1.00 | 1819.66 |
| multieco_nint | 1000 | 6.00 | -3629.22 | -3629.14 | 221.68 | 0.00 | 1.00 | 1819.61 |
| o2range | 1000 | 4.00 | -3622.97 | -3622.93 | 227.89 | 0.00 | 1.00 | 1814.48 |
| btemp | 1000 | 4.00 | -3619.35 | -3619.31 | 231.51 | 0.00 | 1.00 | 1812.67 |
| clownSR | 1000 | 4.00 | -3617.00 | -3616.96 | 233.86 | 0.00 | 1.00 | 1811.50 |
| multibio_int | 1000 | 10.00 | -3601.36 | -3601.14 | 249.68 | 0.00 | 1.00 | 1809.68 |
| surfcurrent | 1000 | 4.00 | -3599.29 | -3599.25 | 251.57 | 0.00 | 1.00 | 1802.65 |
| temp | 1000 | 4.00 | -3598.65 | -3598.61 | 252.21 | 0.00 | 1.00 | 1802.33 |
| host_specificSR | 1000 | 4.00 | -3593.03 | -3592.99 | 257.83 | 0.00 | 1.00 | 1799.52 |
| hostSR | 1000 | 4.00 | -3590.05 | -3590.01 | 260.81 | 0.00 | 1.00 | 1798.02 |
| phyto | 1000 | 4.00 | -3589.96 | -3589.92 | 260.90 | 0.00 | 1.00 | 1797.98 |
| nitrate | 1000 | 4.00 | -3587.89 | -3587.85 | 262.97 | 0.00 | 1.00 | 1796.95 |
| multibio_nint | 1000 | 6.00 | -3587.66 | -3587.57 | 263.25 | 0.00 | 1.00 | 1798.83 |
| chl | 1000 | 4.00 | -3573.64 | -3573.60 | 277.22 | 0.00 | 1.00 | 1789.82 |
| bo2 | 1000 | 4.00 | -3572.51 | -3572.47 | 278.35 | 0.00 | 1.00 | 1789.26 |
| salinity | 1000 | 4.00 | -3570.93 | -3570.88 | 279.94 | 0.00 | 1.00 | 1788.46 |
| sstrange | 1000 | 4.00 | -3570.76 | -3570.72 | 280.10 | 0.00 | 1.00 | 1788.38 |
| tide | 1000 | 4.00 | -3570.76 | -3570.72 | 280.10 | 0.00 | 1.00 | 1788.38 |

**Tab. S9**: Model selection table of Lagrange Multiplier (LM) Spatial Auto-Regressive (SAR) models of *A. clarkii* on inferred deviations from genetic diversity expectations under Isolation-by-distance

| Model | n | K | AIC | AICc | Delta.AICc | AICcWt | Cum.Wt | LogLik |
|---|---|---|---|---|---|---|---|---|
| multiphys_int | 1000 | 10.00 | -3782.52 | -3782.29 | 0.00 | 1.00 | 1.00 | 1900.26 |
| salinity | 1000 | 4.00 | -3694.68 | -3694.64 | 87.65 | 0.00 | 1.00 | 1850.34 |
| multiphys_nint | 1000 | 6.00 | -3693.04 | -3692.95 | 89.34 | 0.00 | 1.00 | 1851.52 |
| mixed_int | 1000 | 130.00 | -3724.87 | -3685.67 | 96.62 | 0.00 | 1.00 | 1991.43 |
| mixed_nint | 1000 | 10.00 | -3678.27 | -3678.05 | 104.25 | 0.00 | 1.00 | 1848.13 |
| multio2d_int | 1000 | 6.00 | -3637.49 | -3637.40 | 144.89 | 0.00 | 1.00 | 1823.74 |
| bo2 | 1000 | 4.00 | -3631.14 | -3631.10 | 151.20 | 0.00 | 1.00 | 1818.57 |
| multio2d_nint | 1000 | 5.00 | -3629.15 | -3629.09 | 153.20 | 0.00 | 1.00 | 1818.58 |
| multitemp_int | 1000 | 10.00 | -3624.92 | -3624.70 | 157.60 | 0.00 | 1.00 | 1821.46 |
| multieco_int | 1000 | 8.00 | -3621.63 | -3621.49 | 160.80 | 0.00 | 1.00 | 1817.82 |
| multitemp_nint | 1000 | 6.00 | -3605.62 | -3605.53 | 176.76 | 0.00 | 1.00 | 1807.81 |
| sstrange | 1000 | 4.00 | -3605.38 | -3605.34 | 176.96 | 0.00 | 1.00 | 1805.69 |
| multibio_int | 1000 | 10.00 | -3584.41 | -3584.18 | 198.11 | 0.00 | 1.00 | 1801.20 |
| hostSR | 1000 | 4.00 | -3581.42 | -3581.38 | 200.92 | 0.00 | 1.00 | 1793.71 |
| host_specificSR | 1000 | 4.00 | -3581.42 | -3581.38 | 200.92 | 0.00 | 1.00 | 1793.71 |
| multieco_nint | 1000 | 5.00 | -3579.54 | -3579.48 | 202.81 | 0.00 | 1.00 | 1793.77 |
| clownSR | 1000 | 4.00 | -3575.56 | -3575.52 | 206.77 | 0.00 | 1.00 | 1790.78 |
| chl | 1000 | 4.00 | -3571.69 | -3571.65 | 210.64 | 0.00 | 1.00 | 1788.85 |
| tide | 1000 | 4.00 | -3568.32 | -3568.28 | 214.01 | 0.00 | 1.00 | 1787.16 |
| multibio_nint | 1000 | 6.00 | -3567.80 | -3567.72 | 214.58 | 0.00 | 1.00 | 1788.90 |
| phyto | 1000 | 4.00 | -3566.63 | -3566.59 | 215.70 | 0.00 | 1.00 | 1786.32 |
| o2range | 1000 | 4.00 | -3565.19 | -3565.15 | 217.15 | 0.00 | 1.00 | 1785.59 |
| nitrate | 1000 | 4.00 | -3564.97 | -3564.92 | 217.37 | 0.00 | 1.00 | 1785.48 |
| multicurrent_nint | 1000 | 6.00 | -3564.76 | -3564.68 | 217.62 | 0.00 | 1.00 | 1787.38 |
| current | 1000 | 4.00 | -3564.53 | -3564.49 | 217.81 | 0.00 | 1.00 | 1785.26 |
| btemp | 1000 | 4.00 | -3564.28 | -3564.24 | 218.06 | 0.00 | 1.00 | 1785.14 |
| temp | 1000 | 4.00 | -3564.27 | -3564.23 | 218.06 | 0.00 | 1.00 | 1785.14 |
| surfcurrent | 1000 | 4.00 | -3564.21 | -3564.17 | 218.12 | 0.00 | 1.00 | 1785.11 |
| multicurrent_int | 1000 | 10.00 | -3561.50 | -3561.28 | 221.01 | 0.00 | 1.00 | 1789.75 |

**Tab. S10**: Model selection table of Lagrange Multiplier (LM) Spatial Auto-Regressive (SAR) models of *A. melanopus* on inferred deviations from genetic diversity expectations under Isolation-by-distance

| Model | n | K | AIC | AICc | Delta.AICc | AICcWt | Cum.Wt | LogLik |
|---|---|---|---|---|---|---|---|---|
| mixed_int | 829 | 130.00 | -5341.16 | -5292.37 | 0.00 | 1.00 | 1.00 | 2799.58 |
| mixed_nint | 829 | 10.00 | -5116.05 | -5115.78 | 176.59 | 0.00 | 1.00 | 2567.02 |
| multio2d_int | 829 | 6.00 | -5017.86 | -5017.75 | 274.61 | 0.00 | 1.00 | 2513.93 |
| multio2d_nint | 829 | 5.00 | -5010.34 | -5010.27 | 282.10 | 0.00 | 1.00 | 2509.17 |
| bo2 | 829 | 4.00 | -5001.14 | -5001.10 | 291.27 | 0.00 | 1.00 | 2503.57 |
| multitemp_int | 829 | 10.00 | -4864.49 | -4864.22 | 428.14 | 0.00 | 1.00 | 2441.25 |
| multiphys_int | 829 | 10.00 | -4862.71 | -4862.44 | 429.93 | 0.00 | 1.00 | 2440.35 |
| salinity | 829 | 4.00 | -4854.77 | -4854.72 | 437.65 | 0.00 | 1.00 | 2430.38 |
| multiphys_nint | 829 | 6.00 | -4853.48 | -4853.38 | 438.98 | 0.00 | 1.00 | 2431.74 |
| multitemp_nint | 829 | 6.00 | -4841.68 | -4841.58 | 450.79 | 0.00 | 1.00 | 2425.84 |
| sstrange | 829 | 4.00 | -4841.53 | -4841.49 | 450.88 | 0.00 | 1.00 | 2423.77 |
| multibio_int | 829 | 10.00 | -4643.20 | -4642.93 | 649.44 | 0.00 | 1.00 | 2330.60 |
| btemp | 829 | 4.00 | -4622.38 | -4622.33 | 670.03 | 0.00 | 1.00 | 2314.19 |
| multicurrent_int | 829 | 10.00 | -4594.02 | -4593.75 | 698.62 | 0.00 | 1.00 | 2306.01 |
| multicurrent_nint | 829 | 6.00 | -4574.88 | -4574.78 | 717.59 | 0.00 | 1.00 | 2292.44 |
| multieco_int | 829 | 10.00 | -4573.16 | -4572.89 | 719.47 | 0.00 | 1.00 | 2295.58 |
| multieco_nint | 829 | 6.00 | -4561.04 | -4560.94 | 731.43 | 0.00 | 1.00 | 2285.52 |
| tide | 829 | 4.00 | -4559.01 | -4558.97 | 733.40 | 0.00 | 1.00 | 2282.51 |
| nitrate | 829 | 4.00 | -4554.76 | -4554.71 | 737.66 | 0.00 | 1.00 | 2280.38 |
| multibio_nint | 829 | 6.00 | -4552.95 | -4552.85 | 739.52 | 0.00 | 1.00 | 2281.47 |
| clownSR | 829 | 4.00 | -4523.95 | -4523.90 | 768.46 | 0.00 | 1.00 | 2264.98 |
| phyto | 829 | 4.00 | -4515.81 | -4515.76 | 776.61 | 0.00 | 1.00 | 2260.90 |
| host_specificSR | 829 | 4.00 | -4513.56 | -4513.51 | 778.85 | 0.00 | 1.00 | 2259.78 |
| surfcurrent | 829 | 4.00 | -4505.12 | -4505.07 | 787.30 | 0.00 | 1.00 | 2255.56 |
| temp | 829 | 4.00 | -4503.07 | -4503.03 | 789.34 | 0.00 | 1.00 | 2254.54 |
| hostSR | 829 | 4.00 | -4493.77 | -4493.72 | 798.65 | 0.00 | 1.00 | 2249.88 |
| o2range | 829 | 4.00 | -4488.06 | -4488.02 | 804.35 | 0.00 | 1.00 | 2247.03 |
| current | 829 | 4.00 | -4486.21 | -4486.16 | 806.20 | 0.00 | 1.00 | 2246.11 |
| chl | 829 | 4.00 | -4486.20 | -4486.15 | 806.21 | 0.00 | 1.00 | 2246.10 |

**Tab. S11**: Model selection table of Lagrange Multiplier (LM) Spatial Auto-Regressive (SAR) models of *A. perideraion* on inferred deviations from genetic diversity expectations under Isolation-by-distance

| Model | n | K | AIC | AICc | Delta.AICc | AICcWt | Cum.Wt | LogLik |
|---|---|---|---|---|---|---|---|---|
| mixed_int | 1000 | 130.00 | -5512.70 | -5473.51 | 0.00 | 1.00 | 1.00 | 2885.35 |
| multiphys_int | 1000 | 10.00 | -5318.86 | -5318.64 | 154.87 | 0.00 | 1.00 | 2668.43 |
| mixed_nint | 1000 | 10.00 | -5250.42 | -5250.20 | 223.31 | 0.00 | 1.00 | 2634.21 |
| multitemp_int | 1000 | 10.00 | -5229.39 | -5229.17 | 244.34 | 0.00 | 1.00 | 2623.70 |
| multio2d_nint | 1000 | 5.00 | -5223.36 | -5223.30 | 250.21 | 0.00 | 1.00 | 2615.68 |
| multio2d_int | 1000 | 6.00 | -5222.41 | -5222.33 | 251.18 | 0.00 | 1.00 | 2616.21 |
| bo2 | 1000 | 4.00 | -5222.12 | -5222.08 | 251.43 | 0.00 | 1.00 | 2614.06 |
| multitemp_nint | 1000 | 6.00 | -5212.93 | -5212.84 | 260.66 | 0.00 | 1.00 | 2611.46 |
| btemp | 1000 | 4.00 | -5195.28 | -5195.24 | 278.26 | 0.00 | 1.00 | 2600.64 |
| multibio_int | 1000 | 10.00 | -5164.52 | -5164.30 | 309.21 | 0.00 | 1.00 | 2591.26 |
| nitrate | 1000 | 4.00 | -5161.86 | -5161.82 | 311.69 | 0.00 | 1.00 | 2583.93 |
| multibio_nint | 1000 | 6.00 | -5158.10 | -5158.01 | 315.49 | 0.00 | 1.00 | 2584.05 |
| multiphys_nint | 1000 | 6.00 | -5154.39 | -5154.31 | 319.20 | 0.00 | 1.00 | 2582.20 |
| sstrange | 1000 | 4.00 | -5147.78 | -5147.74 | 325.76 | 0.00 | 1.00 | 2576.89 |
| temp | 1000 | 4.00 | -5144.79 | -5144.75 | 328.75 | 0.00 | 1.00 | 2575.40 |
| phyto | 1000 | 4.00 | -5141.22 | -5141.18 | 332.33 | 0.00 | 1.00 | 2573.61 |
| multieco_int | 1000 | 10.00 | -5135.58 | -5135.36 | 338.15 | 0.00 | 1.00 | 2576.79 |
| current | 1000 | 4.00 | -5124.02 | -5123.98 | 349.52 | 0.00 | 1.00 | 2565.01 |
| hostSR | 1000 | 4.00 | -5123.38 | -5123.34 | 350.17 | 0.00 | 1.00 | 2564.69 |
| multicurrent_int | 1000 | 10.00 | -5123.36 | -5123.14 | 350.37 | 0.00 | 1.00 | 2570.68 |
| multicurrent_nint | 1000 | 6.00 | -5120.76 | -5120.67 | 352.83 | 0.00 | 1.00 | 2565.38 |
| multieco_nint | 1000 | 6.00 | -5119.76 | -5119.68 | 353.83 | 0.00 | 1.00 | 2564.88 |
| host_specificSR | 1000 | 4.00 | -5119.21 | -5119.17 | 354.33 | 0.00 | 1.00 | 2562.61 |
| salinity | 1000 | 4.00 | -5115.20 | -5115.16 | 358.35 | 0.00 | 1.00 | 2560.60 |
| clownSR | 1000 | 4.00 | -5114.89 | -5114.85 | 358.66 | 0.00 | 1.00 | 2560.45 |
| chl | 1000 | 4.00 | -5113.85 | -5113.81 | 359.69 | 0.00 | 1.00 | 2559.93 |
| o2range | 1000 | 4.00 | -5113.26 | -5113.22 | 360.29 | 0.00 | 1.00 | 2559.63 |
| surfcurrent | 1000 | 4.00 | -5113.24 | -5113.20 | 360.30 | 0.00 | 1.00 | 2559.62 |
| tide | 1000 | 4.00 | -5113.08 | -5113.04 | 360.47 | 0.00 | 1.00 | 2559.54 |

**Tab. S12**: Model selection table of Lagrange Multiplier (LM) Spatial Auto-Regressive (SAR) models of *A. polymnus* on inferred deviations from genetic diversity expectations under Isolation-by-distance

| Model | n | K | AIC | AICc | Delta.AICc | AICcWt | Cum.Wt | LogLik |
|---|---|---|---|---|---|---|---|---|
| mixed_int | 1000 | 130.00 | -5261.58 | -5222.38 | 0.00 | 1.00 | 1.00 | 2759.79 |
| mixed_nint | 1000 | 10.00 | -5045.94 | -5045.71 | 176.67 | 0.00 | 1.00 | 2531.97 |
| multitemp_int | 1000 | 10.00 | -4989.72 | -4989.50 | 232.88 | 0.00 | 1.00 | 2503.86 |
| multitemp_nint | 1000 | 6.00 | -4974.34 | -4974.25 | 248.13 | 0.00 | 1.00 | 2492.17 |
| sstrange | 1000 | 4.00 | -4935.08 | -4935.04 | 287.34 | 0.00 | 1.00 | 2470.54 |
| multicurrent_int | 1000 | 10.00 | -4889.78 | -4889.55 | 332.83 | 0.00 | 1.00 | 2453.89 |
| multieco_int | 1000 | 10.00 | -4872.98 | -4872.76 | 349.63 | 0.00 | 1.00 | 2445.49 |
| multieco_nint | 1000 | 6.00 | -4859.59 | -4859.50 | 362.88 | 0.00 | 1.00 | 2434.79 |
| multicurrent_nint | 1000 | 6.00 | -4856.86 | -4856.78 | 365.61 | 0.00 | 1.00 | 2433.43 |
| hostSR | 1000 | 4.00 | -4854.84 | -4854.80 | 367.58 | 0.00 | 1.00 | 2430.42 |
| tide | 1000 | 4.00 | -4851.84 | -4851.80 | 370.59 | 0.00 | 1.00 | 2428.92 |
| host_specificSR | 1000 | 4.00 | -4846.40 | -4846.36 | 376.03 | 0.00 | 1.00 | 2426.20 |
| temp | 1000 | 4.00 | -4838.55 | -4838.51 | 383.87 | 0.00 | 1.00 | 2422.28 |
| multibio_int | 1000 | 10.00 | -4838.41 | -4838.18 | 384.20 | 0.00 | 1.00 | 2428.20 |
| multiphys_nint | 1000 | 6.00 | -4837.20 | -4837.12 | 385.27 | 0.00 | 1.00 | 2423.60 |
| phyto | 1000 | 4.00 | -4834.76 | -4834.72 | 387.66 | 0.00 | 1.00 | 2420.38 |
| multiphys_int | 1000 | 10.00 | -4834.46 | -4834.24 | 388.15 | 0.00 | 1.00 | 2426.23 |
| multibio_nint | 1000 | 6.00 | -4832.00 | -4831.91 | 390.47 | 0.00 | 1.00 | 2421.00 |
| nitrate | 1000 | 4.00 | -4828.33 | -4828.29 | 394.09 | 0.00 | 1.00 | 2417.17 |
| btemp | 1000 | 4.00 | -4825.86 | -4825.82 | 396.57 | 0.00 | 1.00 | 2415.93 |
| clownSR | 1000 | 4.00 | -4819.97 | -4819.93 | 402.45 | 0.00 | 1.00 | 2412.99 |
| salinity | 1000 | 4.00 | -4816.62 | -4816.58 | 405.80 | 0.00 | 1.00 | 2411.31 |
| chl | 1000 | 4.00 | -4813.45 | -4813.41 | 408.97 | 0.00 | 1.00 | 2409.73 |
| current | 1000 | 4.00 | -4812.25 | -4812.21 | 410.17 | 0.00 | 1.00 | 2409.13 |
| o2range | 1000 | 4.00 | -4807.59 | -4807.55 | 414.84 | 0.00 | 1.00 | 2406.79 |
| bo2 | 1000 | 4.00 | -4807.39 | -4807.35 | 415.04 | 0.00 | 1.00 | 2406.69 |
| multio2d_nint | 1000 | 5.00 | -4806.46 | -4806.40 | 415.98 | 0.00 | 1.00 | 2407.23 |
| surfcurrent | 1000 | 4.00 | -4806.37 | -4806.33 | 416.05 | 0.00 | 1.00 | 2406.18 |
| multio2d_int | 1000 | 6.00 | -4804.74 | -4804.66 | 417.73 | 0.00 | 1.00 | 2407.37 |

**Tab. S13**: Model selection table of Lagrange Multiplier (LM) Spatial Auto-Regressive (SAR) models of *A. sandaracinos* on inferred deviations from genetic diversity expectations under Isolation-by-distance

| Model | n | K | AIC | AICc | Delta.AICc | AICcWt | Cum.Wt | LogLik |
|---|---|---|---|---|---|---|---|---|
| mixed_int | 643 | 130.00 | -3256.31 | -3189.78 | 0.00 | 1.00 | 1.00 | 1757.15 |
| multiphys_nint | 643 | 6.00 | -3152.50 | -3152.37 | 37.42 | 0.00 | 1.00 | 1581.25 |
| multiphys_int | 643 | 10.00 | -3152.44 | -3152.09 | 37.70 | 0.00 | 1.00 | 1585.22 |
| mixed_nint | 643 | 10.00 | -3137.73 | -3137.38 | 52.40 | 0.00 | 1.00 | 1577.86 |
| salinity | 643 | 4.00 | -3116.97 | -3116.90 | 72.88 | 0.00 | 1.00 | 1561.48 |
| multieco_int | 643 | 10.00 | -3101.40 | -3101.05 | 88.73 | 0.00 | 1.00 | 1559.70 |
| multicurrent_nint | 643 | 6.00 | -3069.87 | -3069.74 | 120.05 | 0.00 | 1.00 | 1539.93 |
| multicurrent_int | 643 | 10.00 | -3066.97 | -3066.63 | 123.16 | 0.00 | 1.00 | 1542.49 |
| multitemp_int | 643 | 10.00 | -3059.81 | -3059.47 | 130.32 | 0.00 | 1.00 | 1538.91 |
| tide | 643 | 4.00 | -3059.34 | -3059.27 | 130.51 | 0.00 | 1.00 | 1532.67 |
| multitemp_nint | 643 | 6.00 | -3058.53 | -3058.40 | 131.39 | 0.00 | 1.00 | 1534.26 |
| multieco_nint | 643 | 6.00 | -3054.33 | -3054.20 | 135.58 | 0.00 | 1.00 | 1532.17 |
| bo2 | 643 | 4.00 | -3043.74 | -3043.68 | 146.10 | 0.00 | 1.00 | 1524.87 |
| multio2d_nint | 643 | 5.00 | -3042.61 | -3042.51 | 147.27 | 0.00 | 1.00 | 1525.30 |
| sstrange | 643 | 4.00 | -3041.23 | -3041.17 | 148.62 | 0.00 | 1.00 | 1523.62 |
| multio2d_int | 643 | 6.00 | -3040.65 | -3040.52 | 149.27 | 0.00 | 1.00 | 1525.32 |
| current | 643 | 4.00 | -3040.04 | -3039.97 | 149.81 | 0.00 | 1.00 | 1523.02 |
| hostSR | 643 | 4.00 | -3032.97 | -3032.90 | 156.88 | 0.00 | 1.00 | 1519.48 |
| clownSR | 643 | 4.00 | -3032.15 | -3032.09 | 157.70 | 0.00 | 1.00 | 1519.08 |
| btemp | 643 | 4.00 | -3032.12 | -3032.06 | 157.73 | 0.00 | 1.00 | 1519.06 |
| host_specificSR | 643 | 4.00 | -3026.76 | -3026.69 | 163.09 | 0.00 | 1.00 | 1516.38 |
| surfcurrent | 643 | 4.00 | -3025.34 | -3025.28 | 164.50 | 0.00 | 1.00 | 1515.67 |
| nitrate | 643 | 4.00 | -3024.19 | -3024.13 | 165.66 | 0.00 | 1.00 | 1515.10 |
| phyto | 643 | 4.00 | -3023.75 | -3023.69 | 166.10 | 0.00 | 1.00 | 1514.88 |
| temp | 643 | 4.00 | -3023.36 | -3023.30 | 166.49 | 0.00 | 1.00 | 1514.68 |
| chl | 643 | 4.00 | -3022.40 | -3022.34 | 167.45 | 0.00 | 1.00 | 1514.20 |
| multibio_int | 643 | 10.00 | -3022.60 | -3022.25 | 167.53 | 0.00 | 1.00 | 1520.30 |
| o2range | 643 | 4.00 | -3022.03 | -3021.96 | 167.82 | 0.00 | 1.00 | 1514.01 |
| multibio_nint | 643 | 6.00 | -3021.83 | -3021.70 | 168.09 | 0.00 | 1.00 | 1515.91 |

**Tab. S14**: Model selection table of Lagrange Multiplier (LM) Spatial Auto-Regressive (SAR) models of *A. akallopisos* on inferred deviations from effective migration expectations under a stepping-stone process.

| Model | n | K | AIC | AICc | Delta.AICc | AICcWt | Cum.Wt | LogLik |
|---|---|---|---|---|---|---|---|---|
| mixed_int | 732 | 130.00 | 1638.29 | 1694.96 | 0.00 | 1.00 | 1.00 | -690.14 |
| multiphys_int | 732 | 10.00 | 1718.18 | 1718.49 | 23.53 | 0.00 | 1.00 | -850.09 |
| multieco_int | 732 | 10.00 | 1819.76 | 1820.06 | 125.10 | 0.00 | 1.00 | -900.88 |
| multieco_nint | 732 | 6.00 | 1837.83 | 1837.95 | 142.99 | 0.00 | 1.00 | -913.92 |
| mixed_nint | 732 | 10.00 | 1874.56 | 1874.87 | 179.91 | 0.00 | 1.00 | -928.28 |
| multitemp_int | 732 | 10.00 | 1887.90 | 1888.20 | 193.24 | 0.00 | 1.00 | -934.95 |
| clownSR | 732 | 4.00 | 1909.06 | 1909.11 | 214.16 | 0.00 | 1.00 | -951.53 |
| multibio_int | 732 | 10.00 | 1910.35 | 1910.66 | 215.70 | 0.00 | 1.00 | -946.18 |
| multiphys_nint | 732 | 6.00 | 1912.64 | 1912.75 | 217.80 | 0.00 | 1.00 | -951.32 |
| o2range | 732 | 4.00 | 1917.32 | 1917.38 | 222.42 | 0.00 | 1.00 | -955.66 |
| multitemp_nint | 732 | 6.00 | 1917.89 | 1918.01 | 223.05 | 0.00 | 1.00 | -953.95 |
| multio2d_nint | 732 | 5.00 | 1919.15 | 1919.23 | 224.28 | 0.00 | 1.00 | -955.58 |
| multio2d_int | 732 | 6.00 | 1921.10 | 1921.22 | 226.26 | 0.00 | 1.00 | -955.55 |
| hostSR | 732 | 4.00 | 1923.69 | 1923.74 | 228.79 | 0.00 | 1.00 | -958.84 |
| multibio_nint | 732 | 6.00 | 1924.33 | 1924.45 | 229.49 | 0.00 | 1.00 | -957.17 |
| host_specificSR | 732 | 4.00 | 1925.59 | 1925.64 | 230.69 | 0.00 | 1.00 | -959.79 |
| sstrange | 732 | 4.00 | 1925.97 | 1926.02 | 231.07 | 0.00 | 1.00 | -959.98 |
| multicurrent_nint | 732 | 6.00 | 1928.36 | 1928.48 | 233.52 | 0.00 | 1.00 | -959.18 |
| multicurrent_int | 732 | 10.00 | 1929.92 | 1930.23 | 235.27 | 0.00 | 1.00 | -955.96 |
| current | 732 | 4.00 | 1930.76 | 1930.81 | 235.85 | 0.00 | 1.00 | -962.38 |
| surfcurrent | 732 | 4.00 | 1931.17 | 1931.23 | 236.27 | 0.00 | 1.00 | -962.59 |
| temp | 732 | 4.00 | 1931.98 | 1932.04 | 237.08 | 0.00 | 1.00 | -962.99 |
| phyto | 732 | 4.00 | 1932.55 | 1932.60 | 237.65 | 0.00 | 1.00 | -963.27 |
| bo2 | 732 | 4.00 | 1935.75 | 1935.80 | 240.84 | 0.00 | 1.00 | -964.87 |
| salinity | 732 | 4.00 | 1937.53 | 1937.58 | 242.63 | 0.00 | 1.00 | -965.76 |
| tide | 732 | 4.00 | 1937.81 | 1937.86 | 242.90 | 0.00 | 1.00 | -965.90 |
| nitrate | 732 | 4.00 | 1938.07 | 1938.12 | 243.17 | 0.00 | 1.00 | -966.03 |
| chl | 732 | 4.00 | 1938.41 | 1938.47 | 243.51 | 0.00 | 1.00 | -966.21 |
| btemp | 732 | 4.00 | 1938.63 | 1938.69 | 243.73 | 0.00 | 1.00 | -966.32 |

**Tab. S15**: Model selection table of Lagrange Multiplier (LM) Spatial Auto-Regressive (SAR) models of *A. akindynos* on inferred deviations from effective migration expectations under a stepping-stone process.

| Model | n | K | AIC | AICc | Delta.AICc | AICcWt | Cum.Wt | LogLik |
|---|---|---|---|---|---|---|---|---|
| mixed_int | 1000 | 130.00 | -207.21 | -168.01 | 0.00 | 1.00 | 1.00 | 232.60 |
| mixed_nint | 1000 | 10.00 | 136.92 | 137.14 | 305.15 | 0.00 | 1.00 | -59.46 |
| multio2d_nint | 1000 | 5.00 | 165.22 | 165.28 | 333.30 | 0.00 | 1.00 | -78.61 |
| multio2d_int | 1000 | 6.00 | 166.14 | 166.23 | 334.24 | 0.00 | 1.00 | -78.07 |
| multitemp_int | 1000 | 10.00 | 186.35 | 186.57 | 354.58 | 0.00 | 1.00 | -84.17 |
| multitemp_nint | 1000 | 6.00 | 207.21 | 207.29 | 375.31 | 0.00 | 1.00 | -98.60 |
| multiphys_int | 1000 | 10.00 | 224.68 | 224.90 | 392.91 | 0.00 | 1.00 | -103.34 |
| o2range | 1000 | 4.00 | 237.81 | 237.85 | 405.87 | 0.00 | 1.00 | -115.91 |
| nitrate | 1000 | 4.00 | 243.79 | 243.83 | 411.85 | 0.00 | 1.00 | -118.90 |
| btemp | 1000 | 4.00 | 244.85 | 244.89 | 412.90 | 0.00 | 1.00 | -119.42 |
| multibio_nint | 1000 | 6.00 | 246.01 | 246.10 | 414.11 | 0.00 | 1.00 | -118.01 |
| multibio_int | 1000 | 10.00 | 247.97 | 248.19 | 416.21 | 0.00 | 1.00 | -114.99 |
| multiphys_nint | 1000 | 6.00 | 253.57 | 253.66 | 421.67 | 0.00 | 1.00 | -121.79 |
| temp | 1000 | 4.00 | 257.10 | 257.14 | 425.15 | 0.00 | 1.00 | -125.55 |
| phyto | 1000 | 4.00 | 269.31 | 269.35 | 437.36 | 0.00 | 1.00 | -131.65 |
| multicurrent_int | 1000 | 10.00 | 283.47 | 283.69 | 451.70 | 0.00 | 1.00 | -132.73 |
| multieco_nint | 1000 | 6.00 | 307.01 | 307.09 | 475.11 | 0.00 | 1.00 | -148.51 |
| multicurrent_nint | 1000 | 6.00 | 307.20 | 307.28 | 475.30 | 0.00 | 1.00 | -148.60 |
| hostSR | 1000 | 4.00 | 310.18 | 310.22 | 478.23 | 0.00 | 1.00 | -152.09 |
| multieco_int | 1000 | 10.00 | 313.11 | 313.33 | 481.35 | 0.00 | 1.00 | -147.56 |
| bo2 | 1000 | 4.00 | 326.45 | 326.49 | 494.50 | 0.00 | 1.00 | -160.22 |
| host_specificSR | 1000 | 4.00 | 332.45 | 332.49 | 500.50 | 0.00 | 1.00 | -163.22 |
| current | 1000 | 4.00 | 340.48 | 340.52 | 508.53 | 0.00 | 1.00 | -167.24 |
| tide | 1000 | 4.00 | 360.20 | 360.24 | 528.26 | 0.00 | 1.00 | -177.10 |
| clownSR | 1000 | 4.00 | 362.33 | 362.37 | 530.38 | 0.00 | 1.00 | -178.16 |
| chl | 1000 | 4.00 | 373.63 | 373.67 | 541.69 | 0.00 | 1.00 | -183.82 |
| salinity | 1000 | 4.00 | 408.56 | 408.60 | 576.62 | 0.00 | 1.00 | -201.28 |
| surfcurrent | 1000 | 4.00 | 411.34 | 411.38 | 579.39 | 0.00 | 1.00 | -202.67 |
| sstrange | 1000 | 4.00 | 412.61 | 412.65 | 580.67 | 0.00 | 1.00 | -203.31 |

**Tab. S16**: Model selection table of Lagrange Multiplier (LM) Spatial Auto-Regressive (SAR) models of *A. clarkii* on inferred deviations from effective migration expectations under a stepping-stone process.

| Model | n | K | AIC | AICc | Delta.AICc | AICcWt | Cum.Wt | LogLik |
|---|---|---|---|---|---|---|---|---|
| mixed_int | 1000 | 130.00 | 1845.15 | 1884.34 | 0.00 | 1.00 | 1.00 | -793.57 |
| mixed_nint | 1000 | 10.00 | 2103.53 | 2103.76 | 219.41 | 0.00 | 1.00 | -1042.77 |
| multicurrent_int | 1000 | 10.00 | 2115.16 | 2115.38 | 231.04 | 0.00 | 1.00 | -1048.58 |
| multitemp_int | 1000 | 10.00 | 2122.96 | 2123.18 | 238.84 | 0.00 | 1.00 | -1052.48 |
| multio2d_nint | 1000 | 5.00 | 2148.25 | 2148.31 | 263.96 | 0.00 | 1.00 | -1070.12 |
| multio2d_int | 1000 | 6.00 | 2150.24 | 2150.33 | 265.98 | 0.00 | 1.00 | -1070.12 |
| multicurrent_nint | 1000 | 6.00 | 2154.15 | 2154.23 | 269.89 | 0.00 | 1.00 | -1072.07 |
| multitemp_nint | 1000 | 6.00 | 2154.76 | 2154.84 | 270.50 | 0.00 | 1.00 | -1072.38 |
| bo2 | 1000 | 4.00 | 2157.77 | 2157.81 | 273.47 | 0.00 | 1.00 | -1075.89 |
| sstrange | 1000 | 4.00 | 2158.69 | 2158.73 | 274.39 | 0.00 | 1.00 | -1076.35 |
| tide | 1000 | 4.00 | 2158.72 | 2158.76 | 274.42 | 0.00 | 1.00 | -1076.36 |
| clownSR | 1000 | 4.00 | 2161.09 | 2161.13 | 276.79 | 0.00 | 1.00 | -1077.54 |
| multieco_nint | 1000 | 5.00 | 2163.09 | 2163.15 | 278.81 | 0.00 | 1.00 | -1077.54 |
| multieco_int | 1000 | 8.00 | 2168.31 | 2168.45 | 284.11 | 0.00 | 1.00 | -1077.15 |
| hostSR | 1000 | 4.00 | 2172.55 | 2172.59 | 288.25 | 0.00 | 1.00 | -1083.28 |
| host_specificSR | 1000 | 4.00 | 2172.55 | 2172.59 | 288.25 | 0.00 | 1.00 | -1083.28 |
| multiphys_int | 1000 | 10.00 | 2175.04 | 2175.27 | 290.92 | 0.00 | 1.00 | -1078.52 |
| multiphys_nint | 1000 | 6.00 | 2178.88 | 2178.96 | 294.62 | 0.00 | 1.00 | -1084.44 |
| o2range | 1000 | 4.00 | 2178.99 | 2179.03 | 294.69 | 0.00 | 1.00 | -1086.50 |
| salinity | 1000 | 4.00 | 2180.54 | 2180.58 | 296.24 | 0.00 | 1.00 | -1087.27 |
| multibio_int | 1000 | 10.00 | 2180.47 | 2180.69 | 296.35 | 0.00 | 1.00 | -1081.23 |
| chl | 1000 | 4.00 | 2181.02 | 2181.06 | 296.72 | 0.00 | 1.00 | -1087.51 |
| multibio_nint | 1000 | 6.00 | 2181.48 | 2181.56 | 297.22 | 0.00 | 1.00 | -1085.74 |
| phyto | 1000 | 4.00 | 2182.03 | 2182.07 | 297.73 | 0.00 | 1.00 | -1088.01 |
| current | 1000 | 4.00 | 2182.48 | 2182.52 | 298.18 | 0.00 | 1.00 | -1088.24 |
| surfcurrent | 1000 | 4.00 | 2182.57 | 2182.61 | 298.26 | 0.00 | 1.00 | -1088.28 |
| temp | 1000 | 4.00 | 2183.60 | 2183.64 | 299.30 | 0.00 | 1.00 | -1088.80 |
| btemp | 1000 | 4.00 | 2184.09 | 2184.13 | 299.78 | 0.00 | 1.00 | -1089.04 |
| nitrate | 1000 | 4.00 | 2184.53 | 2184.57 | 300.23 | 0.00 | 1.00 | -1089.27 |

**Tab. S17**: Model selection table of Lagrange Multiplier (LM) Spatial Auto-Regressive (SAR) models of *A. melanopus* on inferred deviations from effective migration expectations under a stepping-stone process.

| Model | n | K | AIC | AICc | Delta.AICc | AICcWt | Cum.Wt | LogLik |
|---|---|---|---|---|---|---|---|---|
| mixed_int | 603 | 130.00 | 968.80 | 1040.96 | 0.00 | 1.00 | 1.00 | -355.40 |
| multiphys_nint | 603 | 6.00 | 1120.10 | 1120.24 | 79.28 | 0.00 | 1.00 | -555.05 |
| multiphys_int | 603 | 10.00 | 1122.72 | 1123.09 | 82.13 | 0.00 | 1.00 | -552.36 |
| multitemp_int | 603 | 10.00 | 1159.31 | 1159.68 | 118.72 | 0.00 | 1.00 | -570.65 |
| salinity | 603 | 4.00 | 1160.65 | 1160.72 | 119.76 | 0.00 | 1.00 | -577.32 |
| multicurrent_int | 603 | 10.00 | 1174.34 | 1174.71 | 133.75 | 0.00 | 1.00 | -578.17 |
| multio2d_int | 603 | 6.00 | 1185.73 | 1185.88 | 144.91 | 0.00 | 1.00 | -587.87 |
| mixed_nint | 603 | 10.00 | 1187.50 | 1187.87 | 146.91 | 0.00 | 1.00 | -584.75 |
| multio2d_nint | 603 | 5.00 | 1192.41 | 1192.51 | 151.55 | 0.00 | 1.00 | -592.20 |
| bo2 | 603 | 4.00 | 1192.88 | 1192.95 | 151.99 | 0.00 | 1.00 | -593.44 |
| multibio_nint | 603 | 6.00 | 1204.56 | 1204.70 | 163.74 | 0.00 | 1.00 | -597.28 |
| temp | 603 | 4.00 | 1205.71 | 1205.78 | 164.82 | 0.00 | 1.00 | -599.86 |
| nitrate | 603 | 4.00 | 1205.86 | 1205.92 | 164.96 | 0.00 | 1.00 | -599.93 |
| chl | 603 | 4.00 | 1207.90 | 1207.97 | 167.01 | 0.00 | 1.00 | -600.95 |
| clownSR | 603 | 4.00 | 1208.36 | 1208.43 | 167.47 | 0.00 | 1.00 | -601.18 |
| o2range | 603 | 4.00 | 1208.88 | 1208.95 | 167.99 | 0.00 | 1.00 | -601.44 |
| current | 603 | 4.00 | 1208.91 | 1208.98 | 168.02 | 0.00 | 1.00 | -601.46 |
| btemp | 603 | 4.00 | 1208.97 | 1209.04 | 168.08 | 0.00 | 1.00 | -601.49 |
| tide | 603 | 4.00 | 1209.30 | 1209.37 | 168.41 | 0.00 | 1.00 | -601.65 |
| multitemp_nint | 603 | 6.00 | 1209.33 | 1209.47 | 168.51 | 0.00 | 1.00 | -599.66 |
| surfcurrent | 603 | 4.00 | 1209.82 | 1209.88 | 168.92 | 0.00 | 1.00 | -601.91 |
| multibio_int | 603 | 10.00 | 1209.54 | 1209.91 | 168.95 | 0.00 | 1.00 | -595.77 |
| multieco_nint | 603 | 6.00 | 1209.83 | 1209.98 | 169.01 | 0.00 | 1.00 | -599.92 |
| multicurrent_nint | 603 | 6.00 | 1210.10 | 1210.24 | 169.28 | 0.00 | 1.00 | -600.05 |
| phyto | 603 | 4.00 | 1210.22 | 1210.29 | 169.33 | 0.00 | 1.00 | -602.11 |
| sstrange | 603 | 4.00 | 1210.47 | 1210.53 | 169.57 | 0.00 | 1.00 | -602.23 |
| host_specificSR | 603 | 4.00 | 1210.63 | 1210.70 | 169.74 | 0.00 | 1.00 | -602.31 |
| hostSR | 603 | 4.00 | 1211.01 | 1211.08 | 170.12 | 0.00 | 1.00 | -602.51 |
| multieco_int | 603 | 10.00 | 1216.58 | 1216.95 | 175.99 | 0.00 | 1.00 | -599.29 |

**Tab. S18**: Model selection table of Lagrange Multiplier (LM) Spatial Auto-Regressive (SAR) models of *A. perideraion* on inferred deviations from effective migration expectations under a stepping-stone process.

| Model | n | K | AIC | AICc | Delta.AICc | AICcWt | Cum.Wt | LogLik |
|---|---|---|---|---|---|---|---|---|
| mixed_int | 537 | 130.00 | 1406.23 | 1490.12 | 0.00 | 1.00 | 1.00 | -574.12 |
| multiphys_int | 537 | 10.00 | 1596.55 | 1596.97 | 106.85 | 0.00 | 1.00 | -789.28 |
| multieco_int | 537 | 10.00 | 1617.16 | 1617.58 | 127.46 | 0.00 | 1.00 | -799.58 |
| mixed_nint | 537 | 10.00 | 1625.79 | 1626.21 | 136.09 | 0.00 | 1.00 | -803.90 |
| multieco_nint | 537 | 6.00 | 1637.45 | 1637.61 | 147.49 | 0.00 | 1.00 | -813.73 |
| multicurrent_int | 537 | 10.00 | 1652.48 | 1652.90 | 162.77 | 0.00 | 1.00 | -817.24 |
| multiphys_nint | 537 | 6.00 | 1657.37 | 1657.53 | 167.41 | 0.00 | 1.00 | -823.69 |
| multitemp_nint | 537 | 6.00 | 1660.04 | 1660.20 | 170.08 | 0.00 | 1.00 | -825.02 |
| clownSR | 537 | 4.00 | 1660.58 | 1660.65 | 170.53 | 0.00 | 1.00 | -827.29 |
| host_specificSR | 537 | 4.00 | 1661.16 | 1661.24 | 171.11 | 0.00 | 1.00 | -827.58 |
| multitemp_int | 537 | 10.00 | 1661.12 | 1661.53 | 171.41 | 0.00 | 1.00 | -821.56 |
| multicurrent_nint | 537 | 6.00 | 1668.67 | 1668.83 | 178.71 | 0.00 | 1.00 | -829.34 |
| salinity | 537 | 4.00 | 1671.66 | 1671.74 | 181.61 | 0.00 | 1.00 | -832.83 |
| btemp | 537 | 4.00 | 1671.92 | 1671.99 | 181.87 | 0.00 | 1.00 | -832.96 |
| sstrange | 537 | 4.00 | 1672.00 | 1672.07 | 181.95 | 0.00 | 1.00 | -833.00 |
| phyto | 537 | 4.00 | 1673.47 | 1673.54 | 183.42 | 0.00 | 1.00 | -833.73 |
| multibio_nint | 537 | 6.00 | 1673.89 | 1674.04 | 183.92 | 0.00 | 1.00 | -831.94 |
| multibio_int | 537 | 10.00 | 1675.51 | 1675.93 | 185.81 | 0.00 | 1.00 | -828.76 |
| nitrate | 537 | 4.00 | 1677.55 | 1677.62 | 187.50 | 0.00 | 1.00 | -835.77 |
| multio2d_int | 537 | 6.00 | 1677.73 | 1677.89 | 187.77 | 0.00 | 1.00 | -833.86 |
| temp | 537 | 4.00 | 1681.89 | 1681.97 | 191.84 | 0.00 | 1.00 | -837.95 |
| tide | 537 | 4.00 | 1682.63 | 1682.70 | 192.58 | 0.00 | 1.00 | -838.31 |
| current | 537 | 4.00 | 1683.62 | 1683.70 | 193.57 | 0.00 | 1.00 | -838.81 |
| hostSR | 537 | 4.00 | 1685.32 | 1685.39 | 195.27 | 0.00 | 1.00 | -839.66 |
| chl | 537 | 4.00 | 1687.36 | 1687.43 | 197.31 | 0.00 | 1.00 | -840.68 |
| bo2 | 537 | 4.00 | 1691.73 | 1691.80 | 201.68 | 0.00 | 1.00 | -842.86 |
| surfcurrent | 537 | 4.00 | 1693.21 | 1693.28 | 203.16 | 0.00 | 1.00 | -843.60 |
| multio2d_nint | 537 | 5.00 | 1693.72 | 1693.83 | 203.71 | 0.00 | 1.00 | -842.86 |
| o2range | 537 | 4.00 | 1696.45 | 1696.52 | 206.40 | 0.00 | 1.00 | -845.22 |

**Tab. S19**: Model selection table of Lagrange Multiplier (LM) Spatial Auto-Regressive (SAR) models of *A. polymnus* on inferred deviations from effective migration expectations under a stepping-stone process.

| Model | n | K | AIC | AICc | Delta.AICc | AICcWt | Cum.Wt | LogLik |
|---|---|---|---|---|---|---|---|---|
| mixed_nint | 764 | 10.00 | 950.46 | 950.75 | 0.00 | 1.00 | 1.00 | -466.23 |
| multiphys_int | 764 | 10.00 | 983.82 | 984.12 | 33.36 | 0.00 | 1.00 | -482.91 |
| mixed_int | 764 | 130.00 | 945.01 | 998.81 | 48.06 | 0.00 | 1.00 | -343.50 |
| salinity | 764 | 4.00 | 1041.58 | 1041.63 | 90.88 | 0.00 | 1.00 | -517.79 |
| multiphys_nint | 764 | 6.00 | 1044.76 | 1044.87 | 94.12 | 0.00 | 1.00 | -517.38 |
| multieco_int | 764 | 10.00 | 1058.30 | 1058.59 | 107.84 | 0.00 | 1.00 | -520.15 |
| multitemp_int | 764 | 10.00 | 1063.75 | 1064.04 | 113.29 | 0.00 | 1.00 | -522.87 |
| hostSR | 764 | 4.00 | 1067.75 | 1067.80 | 117.05 | 0.00 | 1.00 | -530.87 |
| multieco_nint | 764 | 6.00 | 1068.43 | 1068.55 | 117.79 | 0.00 | 1.00 | -529.22 |
| multio2d_nint | 764 | 5.00 | 1087.79 | 1087.87 | 137.12 | 0.00 | 1.00 | -539.89 |
| multio2d_int | 764 | 6.00 | 1088.29 | 1088.40 | 137.65 | 0.00 | 1.00 | -539.15 |
| bo2 | 764 | 4.00 | 1092.88 | 1092.94 | 142.18 | 0.00 | 1.00 | -543.44 |
| phyto | 764 | 4.00 | 1111.87 | 1111.92 | 161.17 | 0.00 | 1.00 | -552.93 |
| multibio_int | 764 | 10.00 | 1112.25 | 1112.54 | 161.79 | 0.00 | 1.00 | -547.12 |
| multibio_nint | 764 | 6.00 | 1115.43 | 1115.54 | 164.79 | 0.00 | 1.00 | -552.72 |
| btemp | 764 | 4.00 | 1121.68 | 1121.74 | 170.98 | 0.00 | 1.00 | -557.84 |
| clownSR | 764 | 4.00 | 1124.82 | 1124.87 | 174.12 | 0.00 | 1.00 | -559.41 |
| multitemp_nint | 764 | 6.00 | 1124.95 | 1125.07 | 174.31 | 0.00 | 1.00 | -557.48 |
| host_specificSR | 764 | 4.00 | 1132.19 | 1132.25 | 181.49 | 0.00 | 1.00 | -563.10 |
| nitrate | 764 | 4.00 | 1135.98 | 1136.04 | 185.28 | 0.00 | 1.00 | -564.99 |
| temp | 764 | 4.00 | 1137.63 | 1137.68 | 186.93 | 0.00 | 1.00 | -565.81 |
| multicurrent_int | 764 | 10.00 | 1139.84 | 1140.13 | 189.38 | 0.00 | 1.00 | -560.92 |
| multicurrent_nint | 764 | 6.00 | 1143.45 | 1143.56 | 192.81 | 0.00 | 1.00 | -566.73 |
| tide | 764 | 4.00 | 1145.91 | 1145.97 | 195.21 | 0.00 | 1.00 | -569.96 |
| chl | 764 | 4.00 | 1150.53 | 1150.58 | 199.83 | 0.00 | 1.00 | -572.27 |
| o2range | 764 | 4.00 | 1153.26 | 1153.31 | 202.56 | 0.00 | 1.00 | -573.63 |
| current | 764 | 4.00 | 1159.24 | 1159.29 | 208.54 | 0.00 | 1.00 | -576.62 |
| sstrange | 764 | 4.00 | 1161.10 | 1161.15 | 210.40 | 0.00 | 1.00 | -577.55 |
| surfcurrent | 764 | 4.00 | 1161.10 | 1161.16 | 210.40 | 0.00 | 1.00 | -577.55 |

**Tab. S20**: Model selection table of Lagrange Multiplier (LM) Spatial Auto-Regressive (SAR) models of *A. chrysopterus* on inferred deviations from effective migration expectations under a stepping-stone process.

| Model | n | K | AIC | AICc | Delta.AICc | AICcWt | Cum.Wt | LogLik |
|---|---|---|---|---|---|---|---|---|
| mixed_int | 1000 | 130.00 | 244.15 | 283.34 | 0.00 | 1.00 | 1.00 | 6.93 |
| mixed_nint | 1000 | 10.00 | 370.65 | 370.88 | 87.53 | 0.00 | 1.00 | -176.33 |
| multicurrent_int | 1000 | 10.00 | 540.30 | 540.52 | 257.18 | 0.00 | 1.00 | -261.15 |
| multio2d_nint | 1000 | 5.00 | 567.30 | 567.36 | 284.02 | 0.00 | 1.00 | -279.65 |
| multio2d_int | 1000 | 6.00 | 568.41 | 568.49 | 285.15 | 0.00 | 1.00 | -279.21 |
| bo2 | 1000 | 4.00 | 579.89 | 579.93 | 296.59 | 0.00 | 1.00 | -286.95 |
| multicurrent_nint | 1000 | 6.00 | 586.02 | 586.11 | 302.77 | 0.00 | 1.00 | -288.01 |
| multiphys_int | 1000 | 10.00 | 588.26 | 588.48 | 305.14 | 0.00 | 1.00 | -285.13 |
| tide | 1000 | 4.00 | 591.06 | 591.10 | 307.76 | 0.00 | 1.00 | -292.53 |
| multibio_int | 1000 | 10.00 | 619.13 | 619.35 | 336.01 | 0.00 | 1.00 | -300.57 |
| multibio_nint | 1000 | 6.00 | 623.69 | 623.77 | 340.43 | 0.00 | 1.00 | -306.84 |
| multiphys_nint | 1000 | 6.00 | 626.63 | 626.71 | 343.37 | 0.00 | 1.00 | -308.31 |
| multitemp_int | 1000 | 10.00 | 639.40 | 639.63 | 356.28 | 0.00 | 1.00 | -310.70 |
| salinity | 1000 | 4.00 | 640.65 | 640.69 | 357.35 | 0.00 | 1.00 | -317.33 |
| multieco_nint | 1000 | 6.00 | 648.58 | 648.66 | 365.32 | 0.00 | 1.00 | -319.29 |
| host_specificSR | 1000 | 4.00 | 648.68 | 648.72 | 365.38 | 0.00 | 1.00 | -321.34 |
| clownSR | 1000 | 4.00 | 650.23 | 650.27 | 366.93 | 0.00 | 1.00 | -322.11 |
| multitemp_nint | 1000 | 6.00 | 652.05 | 652.14 | 368.80 | 0.00 | 1.00 | -321.03 |
| hostSR | 1000 | 4.00 | 654.01 | 654.05 | 370.70 | 0.00 | 1.00 | -324.00 |
| nitrate | 1000 | 4.00 | 654.38 | 654.42 | 371.08 | 0.00 | 1.00 | -324.19 |
| multieco_int | 1000 | 10.00 | 655.39 | 655.61 | 372.27 | 0.00 | 1.00 | -318.69 |
| sstrange | 1000 | 4.00 | 658.13 | 658.17 | 374.82 | 0.00 | 1.00 | -326.06 |
| btemp | 1000 | 4.00 | 666.31 | 666.35 | 383.01 | 0.00 | 1.00 | -330.15 |
| temp | 1000 | 4.00 | 668.46 | 668.50 | 385.16 | 0.00 | 1.00 | -331.23 |
| phyto | 1000 | 4.00 | 668.95 | 668.99 | 385.65 | 0.00 | 1.00 | -331.48 |
| o2range | 1000 | 4.00 | 671.57 | 671.61 | 388.27 | 0.00 | 1.00 | -332.78 |
| current | 1000 | 4.00 | 673.76 | 673.80 | 390.46 | 0.00 | 1.00 | -333.88 |
| surfcurrent | 1000 | 4.00 | 673.80 | 673.84 | 390.50 | 0.00 | 1.00 | -333.90 |
| chl | 1000 | 4.00 | 673.94 | 673.98 | 390.64 | 0.00 | 1.00 | -333.97 |

**Tab. S21**: Model selection table of Lagrange Multiplier (LM) Spatial Auto-Regressive (SAR) models of *A. sandaracinos* on inferred deviations from effective migration expectations under a stepping-stone process.

| Model | n | K | AIC | AICc | Delta.AICc | AICcWt | Cum.Wt | LogLik |
|---|---|---|---|---|---|---|---|---|
| multiphys_int | 560 | 10.00 | 368.85 | 369.25 | 0.00 | 0.98 | 0.98 | -175.42 |
| mixed_int | 560 | 130.00 | 297.76 | 377.15 | 7.90 | 0.02 | 1.00 | -19.88 |
| multitemp_int | 560 | 10.00 | 431.46 | 431.86 | 62.61 | 0.00 | 1.00 | -206.73 |
| salinity | 560 | 4.00 | 438.22 | 438.29 | 69.04 | 0.00 | 1.00 | -216.11 |
| multiphys_nint | 560 | 6.00 | 441.34 | 441.50 | 72.25 | 0.00 | 1.00 | -215.67 |
| mixed_nint | 560 | 10.00 | 451.47 | 451.87 | 82.62 | 0.00 | 1.00 | -216.74 |
| multitemp_nint | 560 | 6.00 | 458.02 | 458.18 | 88.93 | 0.00 | 1.00 | -224.01 |
| multieco_nint | 560 | 6.00 | 477.86 | 478.02 | 108.77 | 0.00 | 1.00 | -233.93 |
| multieco_int | 560 | 10.00 | 481.22 | 481.62 | 112.37 | 0.00 | 1.00 | -231.61 |
| multibio_int | 560 | 10.00 | 482.36 | 482.76 | 113.51 | 0.00 | 1.00 | -232.18 |
| phyto | 560 | 4.00 | 485.70 | 485.77 | 116.52 | 0.00 | 1.00 | -239.85 |
| host_specificSR | 560 | 4.00 | 485.81 | 485.88 | 116.63 | 0.00 | 1.00 | -239.90 |
| btemp | 560 | 4.00 | 486.16 | 486.23 | 116.98 | 0.00 | 1.00 | -240.08 |
| temp | 560 | 4.00 | 487.11 | 487.18 | 117.93 | 0.00 | 1.00 | -240.55 |
| clownSR | 560 | 4.00 | 487.53 | 487.60 | 118.35 | 0.00 | 1.00 | -240.76 |
| multibio_nint | 560 | 6.00 | 489.49 | 489.64 | 120.39 | 0.00 | 1.00 | -239.74 |
| nitrate | 560 | 4.00 | 494.17 | 494.24 | 124.99 | 0.00 | 1.00 | -244.08 |
| sstrange | 560 | 4.00 | 496.21 | 496.28 | 127.03 | 0.00 | 1.00 | -245.11 |
| multicurrent_int | 560 | 10.00 | 498.91 | 499.32 | 130.07 | 0.00 | 1.00 | -240.46 |
| hostSR | 560 | 4.00 | 501.25 | 501.33 | 132.08 | 0.00 | 1.00 | -247.63 |
| chl | 560 | 4.00 | 507.31 | 507.38 | 138.13 | 0.00 | 1.00 | -250.65 |
| current | 560 | 4.00 | 509.76 | 509.84 | 140.59 | 0.00 | 1.00 | -251.88 |
| o2range | 560 | 4.00 | 510.90 | 510.97 | 141.72 | 0.00 | 1.00 | -252.45 |
| tide | 560 | 4.00 | 511.03 | 511.10 | 141.85 | 0.00 | 1.00 | -252.51 |
| surfcurrent | 560 | 4.00 | 511.81 | 511.88 | 142.63 | 0.00 | 1.00 | -252.91 |
| multicurrent_nint | 560 | 6.00 | 511.88 | 512.03 | 142.78 | 0.00 | 1.00 | -250.94 |
| bo2 | 560 | 4.00 | 512.31 | 512.38 | 143.13 | 0.00 | 1.00 | -253.15 |
| multio2d_int | 560 | 6.00 | 512.29 | 512.44 | 143.19 | 0.00 | 1.00 | -251.15 |
| multio2d_nint | 560 | 5.00 | 512.81 | 512.92 | 143.67 | 0.00 | 1.00 | -252.41 |
